# Antibodies that neutralize all current SARS-CoV-2 variants of concern by conformational locking

**DOI:** 10.1101/2023.04.08.536123

**Authors:** Lihong Liu, Ryan G. Casner, Yicheng Guo, Qian Wang, Sho Iketani, Jasper Fuk-Woo Chan, Jian Yu, Bernadeta Dadonaite, Manoj S. Nair, Hiroshi Mohri, Eswar R. Reddem, Shuofeng Yuan, Vincent Kwok-Man Poon, Chris Chung-Sing Chan, Kwok-Yung Yuen, Zizhang Sheng, Yaoxing Huang, Jesse D. Bloom, Lawrence Shapiro, David D. Ho

## Abstract

SARS-CoV-2 continues to evolve and evade most existing neutralizing antibodies, including all clinically authorized antibodies. We have isolated and characterized two human monoclonal antibodies, 12-16 and 12-19, which exhibited neutralizing activities against all SARS-CoV-2 variants tested, including BQ.1.1 and XBB.1.5. They also blocked infection in hamsters challenged with Omicron BA.1 intranasally. Structural analyses revealed both antibodies targeted a conserved quaternary epitope located at the interface between the N-terminal domain and subdomain 1, revealing a previously unrecognized site of vulnerability on SARS-CoV-2 spike. These antibodies prevent viral receptor engagement by locking the receptor-binding domain of spike in the down conformation, revealing a novel mechanism of virus neutralization for non-RBD antibodies. Deep mutational scanning showed that SARS-CoV-2 could mutate to escape 12-19, but the responsible mutations are rarely found in circulating viruses. Antibodies 12-16 and 12-19 hold promise as prophylactic agents for immunocompromised persons who do not respond robustly to COVID-19 vaccines.

## INTRODUCTION

To date coronavirus disease 2019 (COVID-19) caused by severe acute respiratory syndrome coronavirus 2 (SARS-CoV-2) has been confirmed in over 762 million cases, along with over 6.89 million deaths worldwide^1^. To mitigate virus spread and disease impact, interventional measures such as vaccines, antiviral drugs, and monoclonal antibodies (mAbs), have been successfully developed and deployed^2–4^. Numerous studies have shown that immunity acquired through vaccination and/or natural infection can provide robust protection against severe disease, hospitalization, and death, as well as effectively reducing virus transmission^5–8^. However, the emergence of increasingly immune-evasive SARS-CoV-2 Omicron subvariants, along with waning immunity over time, poses significant challenges to the efficacy of vaccines and mAbs^9–19^. In particular, the currently predominant Omicron subvariants BQ.1.1 and XBB.1.5 have acquired many more mutations in their spike glycoprotein, resulting in marked or complete resistance to neutralization by human polyclonal sera, and by the mAb combination known as Evusheld (tixagevimab and cilgavimab)^9, 16, 20^, which had been effective in protecting immunocompromised individuals who did not respond robustly to COVID-19 vaccines. Their need highlights the urgency to develop potent and broad neutralizing antibodies against current and future SARS-CoV-2 strains.

Over the course of the COVID-19 pandemic, thousands of neutralizing mAbs targeting a multitude of spike epitopes have been isolated and characterized. These antibodies largely target the receptor-binding domain (RBD)^21–25^, including a cryptic site that is only revealed when the RBD is in the “up” position^26–29^. A minority of neutralizing antibodies target the N-terminal domain (NTD)^30–33^, as well as the stem helix of S2^34, 35^, subdomain 1 (SD1)^36–38^, and a quaternary site comprised of NTD and subdomain 2 (SD2)^33^. RBD-directed mAbs are generally more potent, and a number have been shown to be clinically effective as therapeutic or prophylactic agents^39–45^. However, the immunodominance of RBD has exerted strong antibody pressure on SARS-CoV-2 evolution, such that all clinically authorized mAbs are now rendered inactive by the latest Omicron subvariants^9–13^. In fact, out of the thousands of mAbs isolated from many laboratories, only a handful have been reported to adequately neutralize the prevailing viruses in the circulation today. S2-directed mAbs retain their neutralization breadth, but their clinical utility is limited by the lack of potency^34, 35, 46^. Only a small number of S1-directed mAbs retain their neutralizing activity against BQ.1.1 and XBB.1.5, or against the recently emerging CH.1.1 and DS.1 from the BA.2.75 sublineage of Omicron^47^. One subset of active mAbs targets the inner face of the RBD (class 1 or 4), exemplified by SA55, BD56-1302, BD56-1854, and BD57-0129^16, 48^, and another set targets SD1, as exemplified by a mouse mAb S3H3^37^ and a human mAb BA.4/5-5^49^. However, it is expected that SARS-CoV-2 will continue to evolve due to the ever-changing selective pressure from serum antibodies in the population. Therefore, we must anticipate the emergence of future variants that will further threaten our already-depleted arsenal of therapeutic antibodies. An effort to restock with mAbs that could broadly neutralize SARS-CoV-2 is warranted.

We now report two genetically related human mAbs, 12-16 and 12-19, that block receptor binding and neutralize all SARS-CoV-2 variants or subvariants tested, with in vitro potency similar to some of the authorized antibodies. They are also protective in vivo against infection by Omicron BA.1 in hamsters. Interestingly, these mAbs do not target RBD but instead target a quaternary epitope formed by NTD and SD1. Antibody binding to this unique site locks the RBD in the “down” position, uncovering a novel mechanism for receptor interference and virus neutralization. Importantly, mutations within this epitope appear to be rare among currently circulating Omicron subvariants, suggesting this spike region is not subject to strong antibody pressure in the population. Antibodies 12-16 and 12-19 are candidates for clinical development.

## RESULTS

### Isolation and characterization of broadly neutralizing monoclonal antibodies against SARS-CoV-2 variants

To isolate mAbs with broadly neutralizing capacity against SARS-CoV-2, we screened a panel of serum samples from convalescing COVID-19 patients against 11 SARS-CoV-2 variants as well as SARS-CoV. Serum from Patient 12 (**Table S1**) demonstrated a high degree of neutralizing activity against all pseudoviruses tested, with the 50% inhibitory dose (ID_50_) titers in serum ranging from 102 to 3076 (**Figure S1A**). We next used the S2P spike trimer of Beta variant (B.1.351) as a probe to sort for antigen-specific memory B cells from the peripheral blood mononuclear cells (PBMCs) of Patient 12, followed by single-cell RNA-sequencing to determine the paired heavy- and light-chain sequences of the antibody made by each B cell. A total of 27 mAbs were isolated, five of which neutralized both D614G and BA.1 (**Figures S1B and S1C**). Among these, 12-16 and 12-19 stood out, exhibiting good neutralizing activity against authentic WA1 and BA.1 viruses (**Figure S1D**), as well as all tested pseudotyped variants of SARS-CoV-2, including the latest prevailing Omicron subvariants of BQ.1.1, XBB.1.5, and CH.1.1 (**Figures 1A and S1E**). Interestingly, while 12-16 and 12-19 did not bind the spike trimer of D614G and B.1.351 by enzyme-linked immunosorbent assay (ELISA) (**Figure 1B**), they strongly bound to the spike trimer expressed on the cell surface (**Figure 1C**) as determined by flow cytometry. Furthermore, a cell-surface competition binding assay revealed that these two mAbs behaved similarly to soluble dimeric human ACE2-Fc (hACE2-Fc) protein in blocking ACE2 binding to the spike protein of D614G expressed on cell surfaces (**Figure 1D**), initially suggesting that the two antibodies may be directed to the RBD.

**Figure 1.**
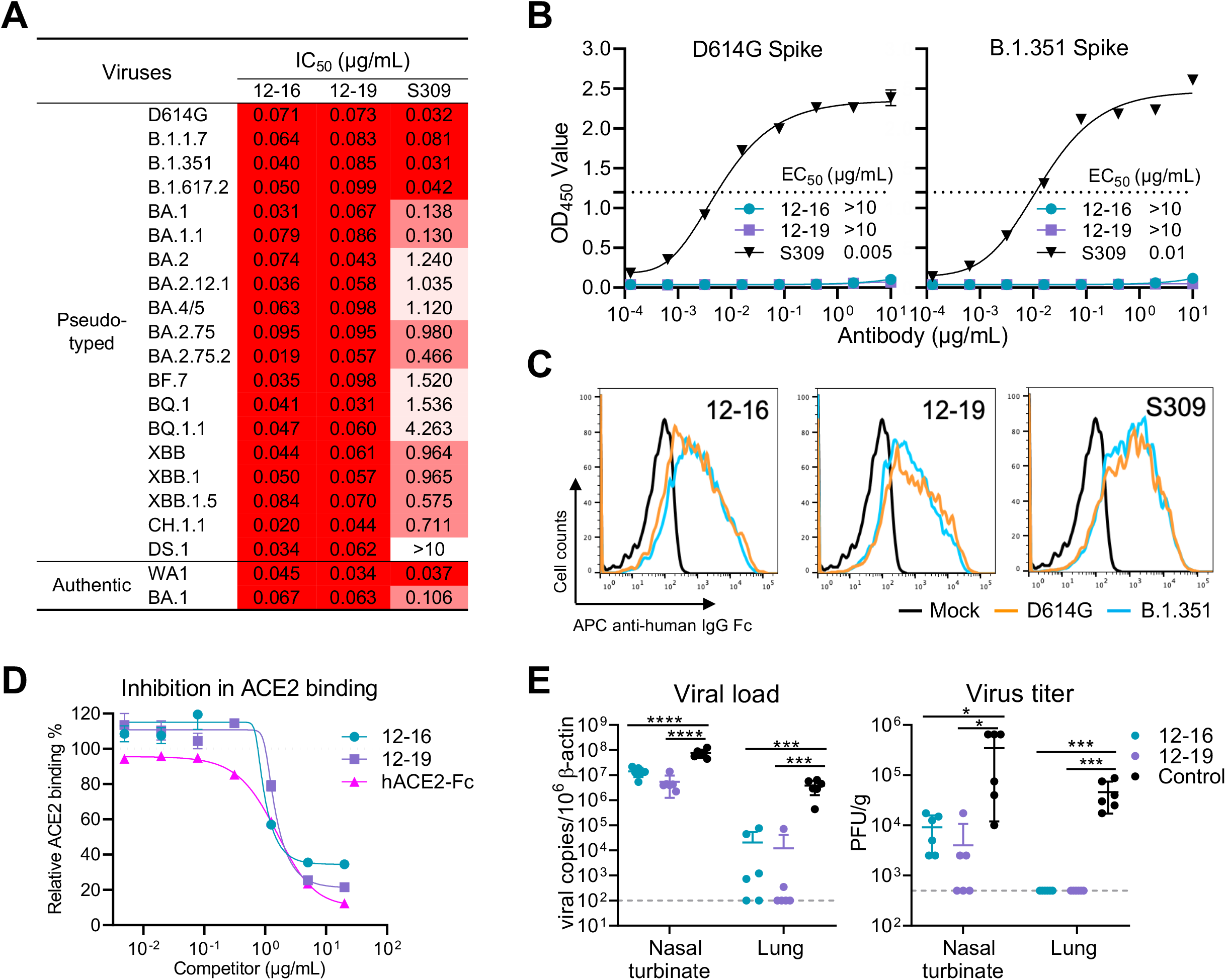
Characterizing the in vitro and in vivo potency and breadth of two neutralizing antibodies. **(A)** Neutralization potency of 12-16 and 12-19 against pseudotyped variants and authentic viruses of SARS-CoV-2. S309 or Sotrovimab was used as a control, which lost potency against Omicron subvariants. **(B)** Binding assay by ELISA indicated that 12-16 and 12-19 could not bind to the SARS-CoV-2 D614G and B.1.351 S2P spike trimers tested. Data are presented as mean ± standard error of the mean (SEM). **(C)** Fluorescence activated cell sorting (FACS) analysis showed that 12-16 and 12-19 well bound to the cell surface-expressed SARS-CoV-2 D614G and B.1.351 spike trimers, indicating they recognize a quaternary epitope on the spike. **(D)** 12-16 and 12-19 inhibited ACE2 binding to cell surface-expressed SARS-CoV-2 D614G spike trimer. hACE2-Fc was used as a positive control. **(E)** Prophylactic efficacy of 12-16 and 12-19 was evaluated in hamsters infected with Omicron variant BA.1. Viral load and titers were measured in trachea and lung 4 days post-infection. Each symbol represents an individual hamster, with a line indicating the mean of each group and error bars indicating the standard deviation. *P* values were determined by unpaired t test. \**p* < 0.05; \*\*\**p* < 0.001; \*\*\*\**p* < 0.0001. Dotted lines indicate assay limits of detection. Each group contained 6 animals. Data in **(A-D)** are representative of those obtained in three independent experiments. See also **Figure S1, S2, Table S1**.

We then evaluated the prophylactic efficacy of 12-16 and 12-19 against Omicron BA.1 in hamsters. Three groups of hamsters (n=6 per group) were administered 10 mg/kg of the indicated mAb via intraperitoneal injection one day before intranasal inoculation with 10^5^ plaque-forming units (PFU) of BA.1. Four days after the virus challenge, nasal turbinate and lung tissues were harvested to quantify SARS-CoV-2. Our results revealed that prophylaxis with either 12-16 or 12-19 significantly reduced the viral RNA copy numbers by approximately 1 log in nasal turbinate and almost 2 logs in lung tissues (**Figure 1E**). Additionally, the administration of 12-16 or 12-19 also reduced the infectious virus titers in both tissues by more than 1 log and down to levels that were no longer detectable by our assay.

Genetically, the heavy chains of 12-16 and 12-19 exhibited high similarity. Specifically, both utilized IGHV3-30*18 and IGHV3-33*01 genes, with the third complementarity determining region of the antibody heavy chain (CDRH3) of 25 and 26 amino acids, respectively (**Figures S2A, S2B, and S2C**). The long CDRH3 of both antibodies were derived from IGHD3-9 and IGHJ6 gene recombination, with four amino acids resulting from N-addition (**Figure S2D**). The light chains of 12-16 and 12-19 were derived from IGLV3-1*01 and IGKV2-29*02, respectively (**Figure S2A**). Both mAbs exhibited low levels of somatic hypermutation, and no rare mutation was detected in either V gene fragments (**Figure S2A, S2B, and S2C**).

### Antibodies 12-16 and 12-19 target a quaternary epitope between NTD and SD1

To determine molecular interactions of 12-16 and 12-19 with spike protein, we employed cryo-EM to visualize Fab fragments of each antibody in complex with S2P-prefusion-stabilized SARS-CoV-2 WA-1 spike protein. Both Fab-spike complexes yielded high-resolution reconstructions (**Figures 2A and 2B**) with global resolutions under 3.1Å (**Figure S3 and Table S2**), allowing for the construction of high-quality molecular models. Surprisingly, rather than recognizing RBD, 12-16 and 12-19 showed similar recognition of a quaternary epitope on the spike situated at the juncture between SD1 and NTD on the side of the spike (**Figures 2A and 2B**). The antibody heavy chains form the primary interactions with SD1, specifically CDR loops 2 and 3, as well as framework region (FR) 3. The epitope on SD1 mainly consists of regions in two loops: the first from residues 557-564 (loop 1), which lies adjacent to NTD, and the second from residues 577-584 (loop 2), which comprises most of the available epitope surface area on the SD1 domain. The epitope on NTD consists of the edge of the NTD β-sandwich (between β sheets 10,13 and 15,16) and the beginning of the N4-loop. The antibody CDRH3 loops serve a critical role, inserting into the crevice between SD1/RBD and NTD. The antibody light chains contact NTD around the N4 loop, an interaction modeled for 12-16, but not for 12-19. Overall, the 12-16 and 12-19 complex structures reveal similar heavy chain-dominated recognition of a new site of vulnerability at the juncture between SD1 and NTD.

**Figure 2.**
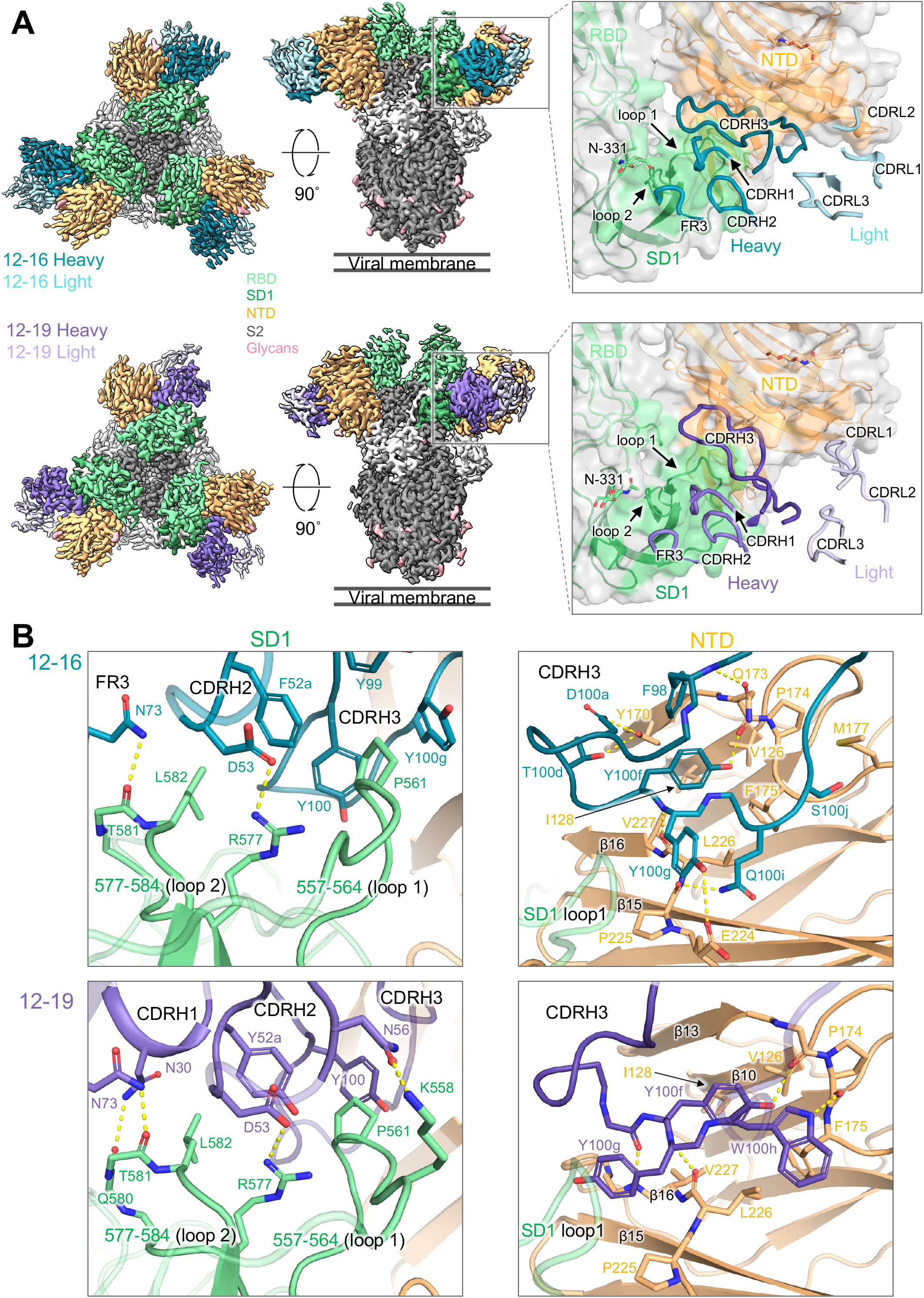
Antibodies 12-16 and 12-19 target a quaternary epitope between SD1 and NTD. **(A)** Cryo-EM reconstructions of antibodies in complex with SARS-CoV-2 D614G spike at resolutions of 3.1 Å for 12-16, and 3.0 Å for 12-19. Each antibody reconstruction is shown first from the top view and second from the side view. The panels show a close-up view of the antibody CDR placement at the quaternary epitope. The orange and green colored surface represent the antibody footprint on NTD and SD1, respectively. **(B)** Interface residues and molecular interactions for 12-16 and 12-19 antibodies showing heavy chain contacts with SD1 (left panel) and CDRH3 contacts with NTD (right panel). The hydrogen bonds are colored in yellow dashed lines. See also **Figure S3 and S4, Table S2**.

Antibodies 12-16 and 12-19 share conserved heavy-chain interactions, including a salt bridge between D53 in the CDRH2 loop and R577 in SD1, as well as hydrogen bonds between N73 in the FR3 region and backbone carbonyls in SD1 loop 2. Furthermore, their CDRH3s are similarly positioned, such that they fill the space between SD1 loop 1 and NTD. In both antibody structures, Y100g is placed between SD1 loop 1 and NTD β16. Y100f is situated nearly identically in both structures, inserting into the NTD β-sandwich and establishing hydrogen bonding interactions. The antibodies bury comparable surface areas in both the SD1 and NTD domains, with most of the buried surface area from the antibody CDRH3 loops, which contain aromatic residues buried in the space between the SD1 loops and between NTD and SD1.

Despite their similarity, the 12-16 and 12-19 Fabs approach the spike protein at somewhat different angles, resulting in differences in their recognition (**Figure S4**). Antibody 12-16 binds higher on spike, with its light chain making more contact with NTD, specifically from contacts of CDR loops 1 and 2 with the NTD N4 loop. The CDR L1 loop forms aliphatic contacts around K182, which reaches towards the CDR L2 loop and forms a salt bridge with D51. In contrast, antibody 12-19 binds slightly lower on the spike and forms additional contacts on SD1 with residues at the beginning of loop 1 (**Figure 2B**). This positioning allows the CDRH3 loop of 12-19 to insert slightly further into the crevice, with the tip of CDRH3 (I100b, L100c) coming closer to RBD and burying surface area with N360, P521, and T523 (**Figure 2B**). This structural difference could possibly explain why 12-19 neutralizes SARS-CoV-2 more completely in vitro than 12-16 does (**Figure S1E**).

### 12-16 and 12-19 neutralize SARS-CoV-2 by locking RBD in the down conformation

How does binding to SD1 and NTD at the side of the spike result in blocking of receptor binding at the top of the spike? Is virus neutralization the consequence of the receptor blockade? To gain insight into these mechanistic questions, we first compared the conformation of SARS-CoV-2 spike protomers in the up-RBD and down-RBD states (**Figure 3A**). We calculated the Cα distance for each of the residues between these two states and found that, as expected, all residues in the RBD region move about 10-50 Å. However, we also observed a substantial (∼10 Å) shift in SD1 and part of the NTD-RBD linker region (**Figure 3B**). In contrast, the NTD and SD2 regions and the S2 subunit showed only minor shifts, with the exception of the ‘FPPR’ segment, as previously reported^50, 51^. Thus, SD1 and part of the NTD-RBD linker region, including the 12-16 and 12-19 epitopes, are significantly affected by the up/down position of RBD.

**Figure 3.**
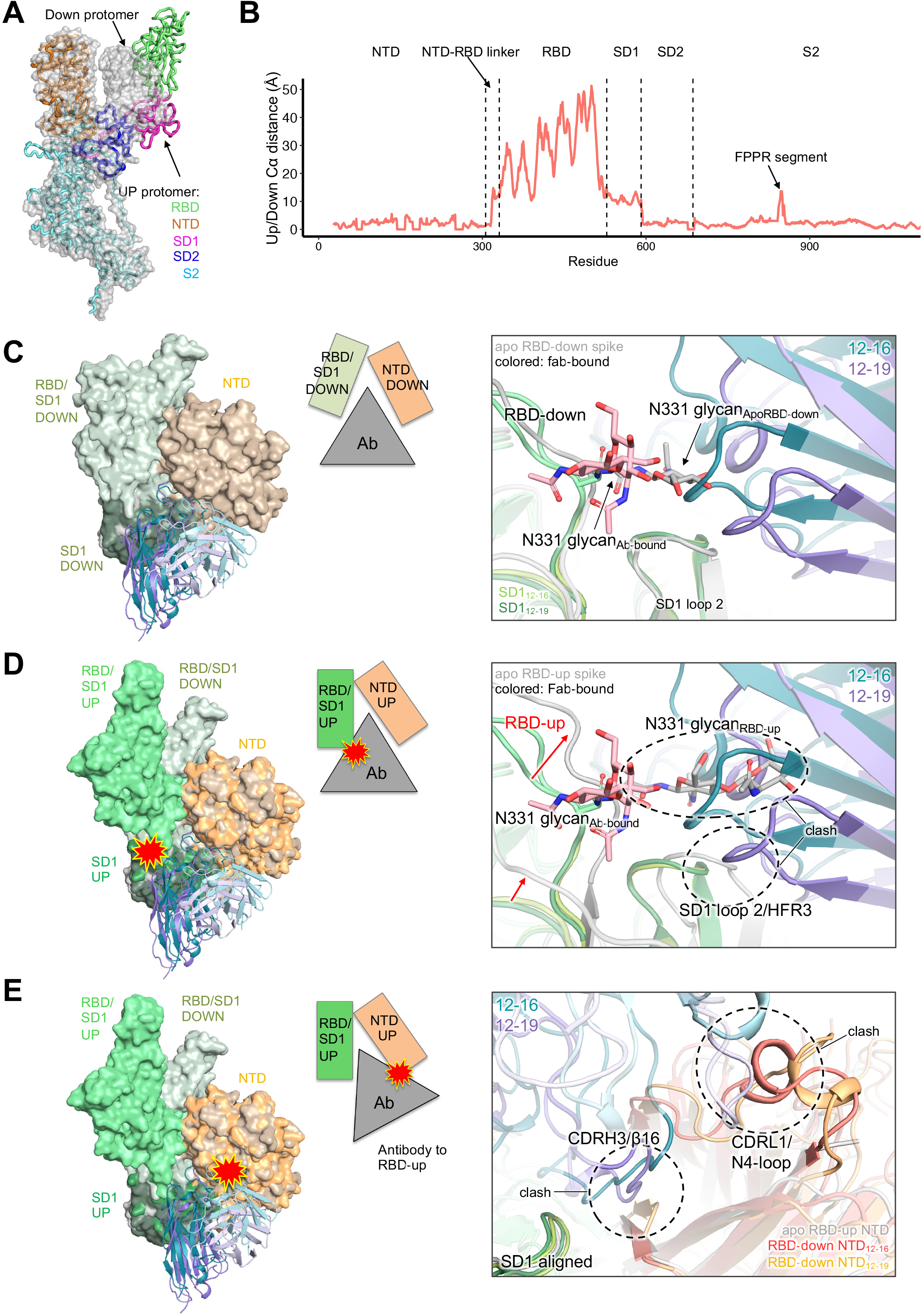
Antibody binding is incompatible with RBD-up state, suggesting the antibody is “locked” in a down state. **(A)** Overlay of an RBD-down (surface) and RBD-up (ribbons) SARS-CoV-2 spike structure (PDB: 7KRR). **(B)** The Cα distance for each residue in the spike between the RBD-up and RBD-down state. **(C)** The 12-16 and 12-19 antibody structures (colored) were superimposed onto an apo SARS-CoV-2 spike structure (pdb: 6XM5) with RBD-down (gray). When the RBD is down and the antibody is bound, the N331 glycan moves out of the way and is nicely accommodated next to the antibody heavy chain FR3 region. **(D)** The 12-16 and 12-19 antibody structures (colored) were superimposed onto an apo SARS-CoV-2 spike structure (pdb: 6XM3) with RBD-up (grey). When the RBD moves up, there are clashes between the N331 glycan and the heavy chain and the RBD strand leading to SD1 (residue ∼530) and heavy chain FR3. **(E)** The 12-16 and 12-19 antibody structures (colored) was superimposed onto an apo SARS-CoV-2 spike structure with RBD-up (gray) using an alignment of the SD1 region of the epitope (residues 531-588) to simulate an “RBD-up bound antibody”. The “RBD-up bound antibody” would clash with the original position of NTD (panel C). In this case, the CDRH3 would clash with the β16 sheet of NTD, and the CDRL1 would clash with the N4-loop. Compared to the position of the NTD in the apo RBD-up structure, the antibody CDRH3 would clash with the β16 sheet of NTD, but it is unclear if the N4-loop would clash. See also **Figure S5**, **Movie 1**, and **Movie 2**.

We next performed particle classification to reveal the approximate number of Fabs bound per spike (**Figure S5**). For 12-16, nearly all (96%) of the spike particles were classified into 3 Fabs bound per spike, whereas for 12-19, most (65%) of the particles were also 3-Fab bound, with the remainder classified into a 2-Fab bound class. Notably, even in the 2-Fab bound class reconstruction, strong density was observed for the 12-19 CDRH3, but density for the outer parts of the Fabs were missing. Further classification and 3D variability analysis in cryoSPARC indicated that the 3-Fab bound spike complexes were very stable, with hardly any flexibility (e.g., in the RBDs) or variability observed (**Movie 1**). For the 2-Fab-bound 12-19 class, the particles were further classified into 3-RBD down and 2-RBD down states, and the motion between RBD up and down states was captured by 3D variability analysis (**Movie 2**) and heterogeneous refinement. Overall, the 3-Fab bound structures were stable without significant variability, but the 2-Fab bound class showed up-and-down mobility of the one RBD that does not have a Fab bound to its neighboring SD1/NTD.

Interestingly, when the spike particles in both datasets were subjected to 3D classification, nearly all the spike particles were observed to be in a 3-RBD down state. This deviates from apo spike structures, where a mix of 1-RBD-up/2-RBD-down and 3-RBD-down is observed^50, 52^. This suggests that the presence of the antibody prevents the observation of a 1-RBD up state, possibly due to the antibody binding locking the spike in a 3-RBD-down conformation. We investigated this hypothesis by structurally analyzing whether 12-16 and 12-19 binding was incompatible with the movement of RBD to an up state. When the antibody models were superimposed onto an unbound spike model with an RBD-down protomer (pdb: 6XM5) (**Figure 3C**), the models aligned closely with minor differences in SD1. However, when the antibody models were superimposed onto an unbound spike model with an RBD-up protomer (pdb: 6XM3) (**Figure 3D**), several incompatibilities with antibody binding were observed. The upward motion of RBD produced clashes between the N331 glycan and the heavy chain, as well as between the RBD strand leading to SD1 (residue ∼530) and heavy chain FR3. Importantly, when the RBD moves up, SD1 swings down and closer to NTD (**Movie 2**), which would be prevented by the placement of the antibody CDRH2 and CDRH3. Additionally, if the antibody were to remain bound to SD1 as SD1 moved into the RBD-up position, the antibody would clash with NTD at two sites: CDRH3 with β16 and CDRL1 with the N4-loop (**Figure 3E**). Thus, the antibody and its CDRH3 forms a ‘wedge’ between SD1 and NTD that prevents the conformational rearrangement necessary for RBD to reach the up conformation. With 3 RBDs locked in the down conformation, the spike cannot engage host receptor ACE2 and gain entry for infection, suggesting that this may be the mechanism of neutralization for 12-16 and 12-19.

### 12-16 and 12-19 block ACE2 binding, as well as ACE2- and CB6-induced S1 shedding

We have shown through our structural analysis (**Figure 3**) that 12-16 and 12-19 neutralize SARS-CoV-2 variants by locking the RBDs in the “down” conformation. As ACE2 only binds to the RBD in the “up” position (**Figure 4A**), it is reasonable to expect that ACE2 binding would be impeded or blocked in the presence of 12-16 and 12-19. To confirm this hypothesis, we conducted a series of biochemistry experiments. First, we examined ACE2 binding to cell surface-expressed SARS-CoV-2 D614G spike in the presence of various competitors targeting a range of epitopes. Along with 12-16 and 12-19, we included the following controls: hACE2-Fc, two ACE2-competing RBD-directed mAbs (2-15 and REGN10987)^21, 39^, two neutralizing NTD supersite-directed mAbs (4-8 and 4-18)^21, 30^, one neutralizing NTD alternative site-directed mAb (5-7)^21, 53^, one non-neutralizing NTD-directed mAb (4-33)^21^, and one non-ACE2-competing S2-directed mAb (S2P6)^34^. The structures for each of these mAbs in complex with spike had previously been solved, with the exception of 4-33. Therefore, we solved the structure of 4-33 in complex with the D614G spike and found that it binds outside of the NTD supersite (**Figure 4A and Table S2**). The overlaid binding epitopes for ACE2 and all of the mAbs are shown in **Figure 4A**. In the competition assay, mAbs 2-15 and REGN10987 were able to compete with ACE2 binding potently, whereas S2P6, 5-7, and 4-33 did not (**Figure 4B**). Interestingly, even though 4-8, 4-18, 12-16, and 12-19 are directed to the NTD region and located distally from the ACE2 binding site, they exhibited discernible to strong competition for ACE2 binding to cell surface expressed D614G spike.

**Figure 4.**
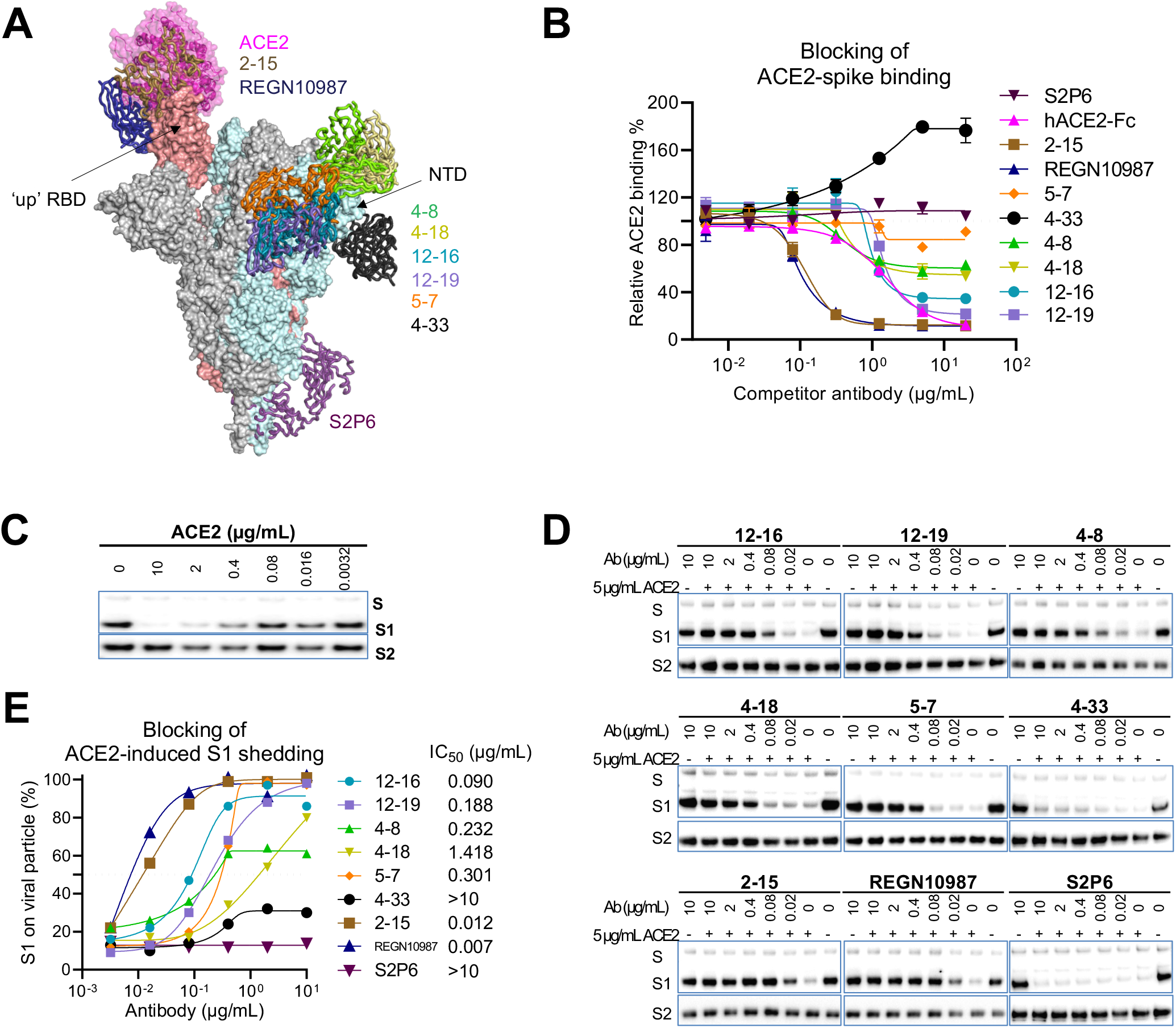
12-16 and 12-19 inhibit ACE2-spike binding and ACE2-induced S1 shedding. **(A)** Human ACE2 and antibodies in complex with SARS-CoV-2 spike with one RBD in the up position (PDB: 7KRR). All of these antibodies can neutralize SARS-CoV-2 except 4-33. **(B)** Competition assay of ACE2 binding to cell surface-expressed SARS-CoV-2 D614G spike in the presence of competitors. The data are shown as the mean ± SEM. **(C)** ACE2-induced S1 shedding from SARS-CoV-2 virions. D614G pseudovirus particles were incubated with hACE2-Fc at different doses for one hour at 37°C before the retained S1 and S2 subunits were determined by western blot. **(D)** Inhibition of ACE2-induced S1 shedding from the spike trimers on SARS-CoV-2 virions by the indicated antibodies. D614G pseudovirus particles were incubated with the indicated antibodies for one hour prior to incubating with 5 µg/mL hACE2-Fc for another one hour. The retained S1 and S2 subunits were determined by western blot. **(E)** The intensities of the S1 and S2 glycoprotein bands in **(D)** were measured and the S1/S2 ratios are shown. Numbers denote the concentration of each antibody which inhibited half of the shedding of S1. The results in **B**, **C, D, and E** are representative of those obtained in two independent experiments. See also **Figure S6**.

Second, as ACE2 binding to spike induces shedding of the S1 glycoprotein^54, 55^ (**Figure 4C**), we further tested if 12-16 and12-19 could block ACE2-induced S1 shedding from viral particles. We tested this by treating SARS-CoV-2 D614G pseudovirus particles with the antibodies before incubating them with hACE2-Fc, and then determining the levels of S1 and S2 glycoproteins on the virions using western blot analysis (**Figures 4D and 4E**). In agreement with the ACE2 competition assay, S2P6 and 4-33 could not block ACE2-induced S1 shedding from virions, whereas RBD-specific mAbs 2-15 and REGN10987 showed the strongest blocking activity. 12-16, 12-19, and the NTD-directed neutralizing antibodies 4-8, 4-18, and 5-7 also protected S1 from ACE2-induced shedding from virions, with efficiencies concordant to their ACE2 competition profiles (**Figure 4B**).

Third, to further corroborate our findings that RBD could not move up in the presence of 12-16 or 12-19, we conducted a similar competition binding assay using CB6, a RBD class 1 neutralizing mAb recognizing RBD only when it is in the up position^41^ (**Figures S6A and S6B**). CB6 itself and the other two RBD-directed mAbs blocked CB6 binding to cell surface-expressed D614G spike, but S2P6, 4-33, and 5-7 did not, as expected. We observed that 12-16 and 12-19 significantly reduced CB6 binding to the spike, while 4-8 and 4-18 showed only minor effects at the two highest doses. Like ACE2, CB6 also triggered shedding of the S1 subunit from the viral particles after binding to D614G spikes (**Figure S6C**). We examined the ability of these mAbs to inhibit CB6-induced S1 shedding and found that only 12-16 and 12-19, but not 4-18, 5-7, 4-33, and S2P6, could block this process (**Figures S6D and S6E**). These results further confirm that 12-16 and 12-19 prevents RBD to be in the up position, most likely from the conformational locking mechanism described in the structural analyses above.

### Neutralization mechanism of 12-16 and 12-19

Our observations that 12-16 and 12-19 could impede S1 shedding mediated by both ACE2 and CB6 (**Figures 4 and S6**), whereas the NTD supersite-directed antibodies like 4-18 were unable to do so, suggest that these two groups of antibodies utilize distinct neutralization mechanisms. Most NTD-directed neutralizing antibodies, including 4-18^30^, target the same antigenic supersite. Previous studies suggest that their neutralization may rely on steric hindrance caused by the Fc^32, 56^. Our structural analysis similarly revealed that the Fc domains of these NTD-supersite mAbs might clash with dimeric ACE2, resulting in neutralization (**Figure S7A**).

We therefore investigated whether the neutralization mechanism of 12-16 and 12-19 involves the steric hindrance arising from the Fc region. To assess this question, we measured the ACE2 binding affinity and neutralization potency of 12-16, 12-19, and 4-18 in the F(ab’)2 and Fab formats, both of which lack Fc (**Figure S7B**). If the Fc region were involved in neutralization, we would expect a decrease in neutralizing potency when Fc is removed while the binding affinity would remain unaffected. Indeed, our data showed that the F(ab’)2 of 12-16, 12-19, and 4-18 had similar binding affinity to spike as their full-length IgG form, while the Fab counterparts showed a reduction in affinity as expected due to the loss in avidity (**Figure 5A** and **S7C**). We then tested their neutralizing activities, finding that while the F(ab’)2 of 12-16 and 12-19 had similar potency as full-length IgG, the neutralization activity of 4-18 was significantly reduced by 10-fold in the F(ab’)2 format compared to the IgG form (**Figure 5B**). As expected from the loss in affinity, the Fab format for all three mAbs tested showed a significant loss in neutralizing activity (**Figure 5B**). The improved binding and neutralization of IgG and F(ab’)2 over the Fab format are likely due to avidity effects. The 12-16 and 12-19 angle of approach on spike would be compatible with spike crosslinking on the virus particle (**Figure 5C**). Collectively, these results showed that NTD supersite-directed mAbs, such as 4-18, could compete with ACE2 through steric hindrance, while NTD-SD1-directed mAbs do not rely on their Fc region for neutralization. Instead, just the Fab region of 12-16 or 12-19 is sufficient to neutralize SARS-CoV-2, as one might expect for the proposed conformational-locking mechanism (**Figure 5C**).

**Figure 5.**
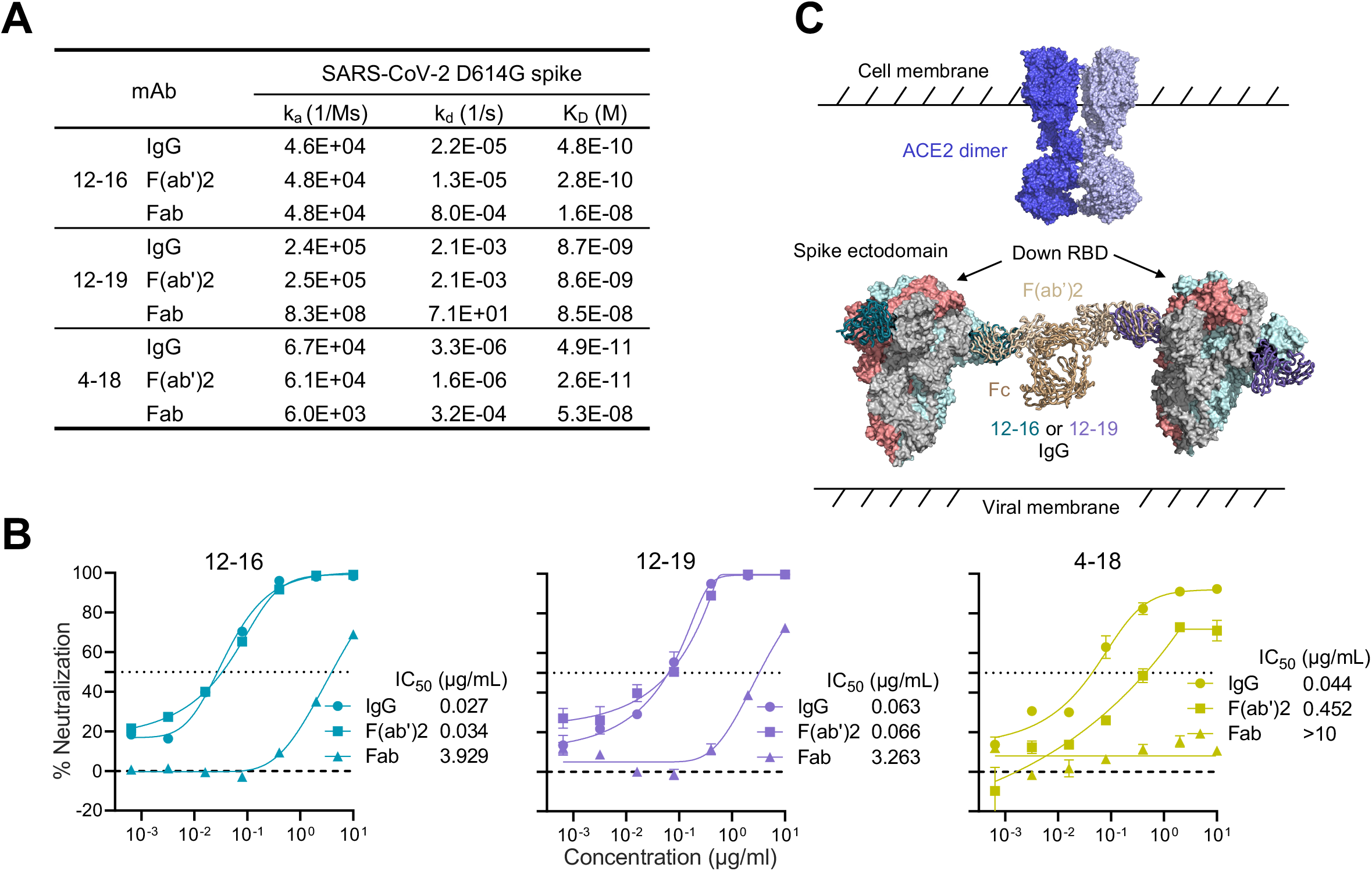
Neutralization of 12-16 and 12-19 is Fc independent. **(A)** Binding kinetics and affinities of 12-16, 12-19, and 4-18 in the formats of IgG, F(ab’)2, and Fab to SARS-CoV-2 D614G S2P spike trimer. **(B)** Neutralization curves and IC50 values of 12-16, 12-19, and 4-18 in the formats of IgG, F(ab’)2, and Fab against D614G. The NTD-directed mAb 4-18 in F(ab’)2 format showed weakened neutralization activity against D614G due to the lack of Fc, while F(ab’)2 of 12-16 and 12-19 retained unimpaired potency, suggesting that the neutralizing activities of 12-16 and 12-19 are Fc independent. Data are shown as mean ± SEM. **(C)** 12-16 and 12-19 maintain the spike at a conformational state with all three RBD in the down position, thereby blocking ACE2 binding. IgG and F(ab’)2 formats would benefit from avidity effects such as spike crosslinking, demonstrated by superimposing 12-16 and 12-19 Fab structures on IgG structure (pdb: 1igt). Data in **A** and **B** are representative from one of three independent experiments. See also **Figure S7**.

### The epitope of antibody 12-19 is highly conserved

Antibodies 12-16 and 12-19 effectively neutralized all the Omicron subvariants tested, including BQ.1.1, XBB.1.5, CH.1.1, and DS.1, with similar potencies as the ancestral D614G strain (**Figure 1A**). To understand 12-19 mutational tolerance, we employed lentivirus-based deep mutational scanning libraries^57^ to map the escape mutations in the background of BA.1 for 12-19. The key spike alterations for escape from this mAb mapped to the base of the N4 loop in NTD (residues 172-176) (**Figure 6A and S8A**). In addition, deletions in the tip of the ß-sheet upstream of the N4 loop and residues at the base of other NTD loops (e.g., 103 and 121) also led to antibody escape. Notably, escape was also observed with mutations in SD1 at the base of the RBD and adjacent to the NTD at residues 522, 561 and 577 (**Figures 6A and S8B**).

**Figure 6.**
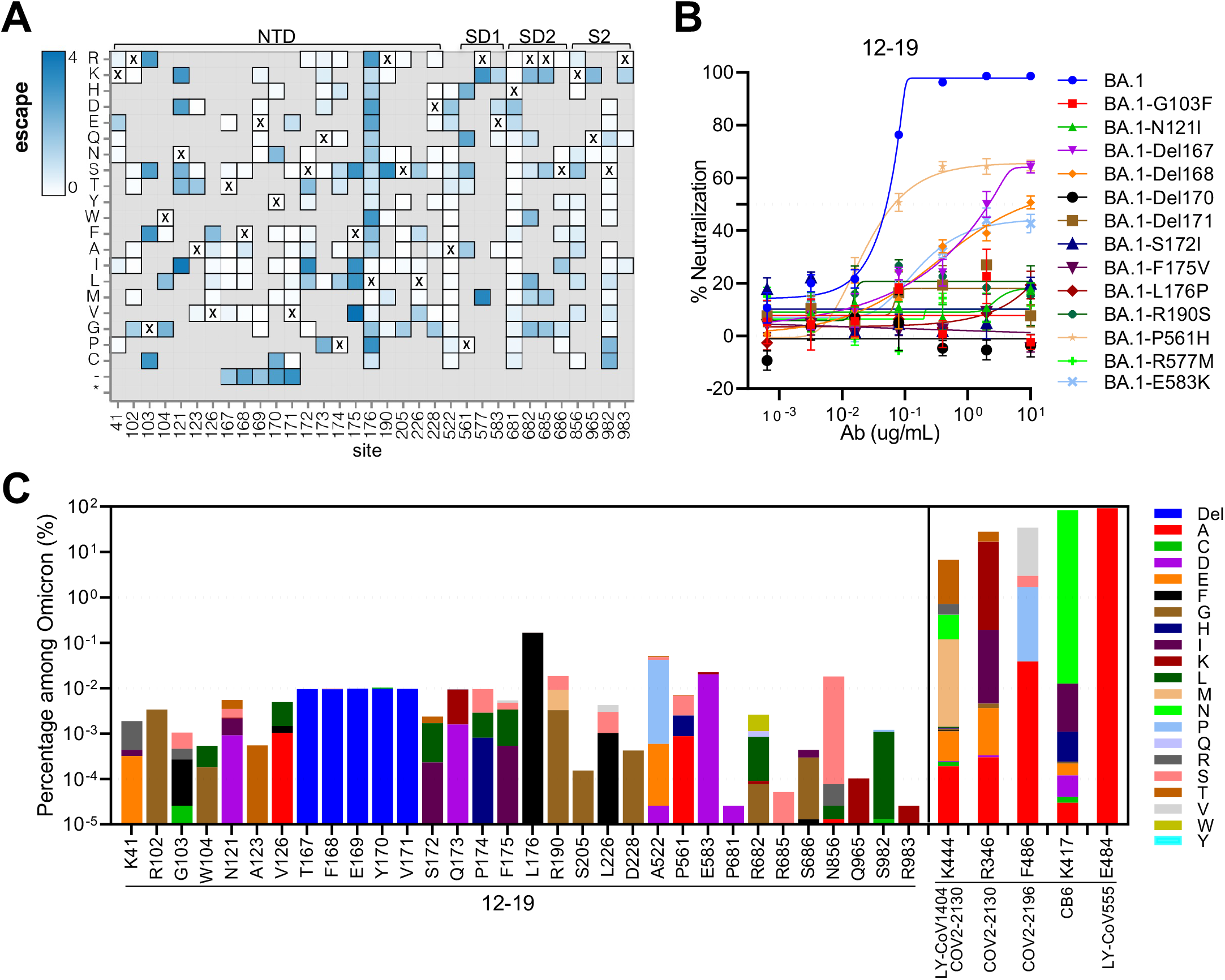
12-19 recognizes a highly conserved epitope. **(A)** Heatmap of mutation escape scores at key sites. Residues marked with X are the wild-type amino acids in BA.1. Amino acids do not present in deep mutational scanning libraries are shown in gray. **(B)** Neutralization resistance to 12-19 conferred by the individual mutations identified by deep mutational scanning in panel **A**. The key mutations were introduced into BA.1 and tested for their sensitivity to 12-19 neutralization. **(C)** Proportion of the key escape mutations of 12-19 in Omicron subvariants, in comparison of the major escape mutations of clinical antibodies. The proportion of escape mutations was determined by analyzing all sequences in GISAID from Omicron BA.1 using the sequence set available as of March 18^th^ 2023. Site numbering is based on the Wuhan-Hu-1 sequence. See https://dms-vep.github.io/SARS-CoV-2_Omicron_BA.1_spike_DMS_12-19/mAb_12-19_escape_plot.html for an interactive version of the heatmap that allows examination of escape across all sites in spike. See also **Figure S8.**

The phenotype of these 12-19 escape mutations were individually validated by introducing each into Omicron BA.1 spike followed by evaluating their neutralization sensitivity in vitro. Each of the 13 introduced mutations substantially or completely impaired the neutralizing activity of 12-19 relative to BA.1 (**Figure 6B**), thus confirming the results from deep mutational scanning. Providentially, these escape mutations are relatively infrequent in the currently circulating Omicron subvariants, with percentages of well below 0.1%, except for L176F whose frequency is 0.2% (**Figure 6C**). These mutational frequencies are generally a thousand-fold lower than those of mutations that escape existing mAbs, including clinically authorized antibodies such as LY-CoV1404 (bebtelovimab), COV2-2130 (cilgavimab), COV2-2196 (tixagevimab), CB6 (etesevimab), and LY-CoV555 (bamlanivimab). To date, the antigenic pressure on this epitope seems to be low, and the alterations in Omicron variants are distant from the binding sites of 12-16 and 12-19 (**Figure S8C**).

## DISCUSSION

Since the onset of the COVID-19 pandemic, SARS-CoV-2 has continuously evolved. The emergence of the Omicron variant and its subsequent subvariants, which possess an unprecedented number of mutations in the spike protein, has resulted in significant resistance to sera obtained from infected and vaccinated individuals, as well as to almost all reported SARS-CoV-2 mAbs^9–19^. Furthermore, recent reports indicate that both the BQ and XBB sublineages are entirely resistant to a mAb combination known as Evusheld, which had been authorized for the prevention of COVID-19. This loss leaves no authorized antibody for clinical use^9, 16, 20^ and presents a significant challenge for the millions of immunocompromised individuals worldwide who do not respond robustly to COVID-19 vaccines. That such persons live in constant fear highlights the urgent need to develop active mAbs for their use as passive immunization. In the current study, we isolated and characterized two genetically related and broadly neutralizing mAbs, 12-16 and 12-19, which retained neutralization activity against all SARS-CoV-2 strains tested, including BQ.1.1, XBB.1.5, and CH.1.1 (**Figures 1A, S1D, and S1E**). Their neutralization potencies in vitro are similar to that of S309 (sotrovimab), which had been an authorized antibody therapeutic. Each of these mAbs also substantially protected hamsters from intranasal challenge with Omicron BA.1 (**Figure 1E**). These features of 12-16 and 12-19 suggest they are promising candidates for clinical development.

We have previously identified an RBD-directed mAb, 2-43, which blocks ACE2 binding and neutralizes SARS-CoV-2 but fails to bind the spike protein in ELISA^21^. At first glance, 12-16 and 12-19 appeared to be similar (**Figures 1A-D**), but such a notion was quickly dispelled when the reconstructions from the cryoEM data showed that these two mAbs recognize quaternary epitopes comprised of NTD and SD1 (**Figure 2**). To the best of our knowledge, these mAbs represent the first report of antibodies targeting both NTD and SD1, revealing a novel site of vulnerability on the SARS-CoV-2 spike. This discovery of a previously unrecognized epitope for broad neutralization suggests that there may still be other conserved sites on SARS-CoV-2 spike to be discovered. Antibodies to such targets may contribute to the residual serum neutralizing activity against latest Omicron sublineages observed among vaccinated individuals, particularly those who have received booster doses^58–61^. Continued identification and understanding of broadly neutralizing mAbs could facilitate the development of future SARS-CoV-2 vaccines.

Mechanistically, it remained unclear why binding to an epitope on the side of spike would lead to interference with receptor binding at the top of the spike. We set out to elucidate this matter by conducting additional structural and biochemical analyses. Particle classification and 3D-variability analysis provided a clue, as antibody binding was observed exclusively in the RBD-down state (**Figure S5**). Structural modeling demonstrated that 12-16 and 12-19 binding to the RBD-up state was incompatible with the upward movement of RBD, implying that upon antibody binding the RBD was “locked” in a down position (**Figure 3**, **Movie 1 and 2**). As this upward movement is required for ACE2 binding to spike, the natural corollary of this “locking” mechanism is that ACE2 binding would be impeded. We confirmed this hypothesis by observing the loss of ACE2 binding and ACE2-induced S1 shedding in the presence of 12-16 and 12-19 (**Figure 4**). Analogous experiments examining the interference of CB6 binding to spike, which also requires an RBD-up state, further confirmed that 12-16 and 12-19 induce a “locked” conformation of spike in a down position (**Figure S6**). Finally, to further demonstrate that it was through this “locking” mechanism that these antibodies neutralized SARS-CoV-2, we showed that the Fc domain was dispensable for the activity of 12-16 and 12-19, indicating that Fc-mediated steric clashes were not required, as would be expected if the “locking” was their main route of neutralization (**Figure 5**). In contrast, we note that an NTD-supersite antibody, 4-18^21, 30^, displayed significant loss of activity without its Fc domain, suggesting that despite the distality of NTD antibodies, they may neutralize through Fc-mediated clashes with ACE2-binding. Collectively, these structural and biochemical studies reveal a unique mechanism of action for 12-16 and 12-19, and in turn extend our understanding of antibody neutralization of SARS-CoV-2.

The results from the pseudovirus-based mutational scanning experiment suggest that 12-19-escape mutations on the spike are not yet prevalent, indicating that its epitope is still relatively conserved and not subject to an inordinate amount of antibody pressure to date (**Figure 6**). These findings suggest that 12-16 and 12-19 may accommodate additional mutations that the virus may acquire in the future, perhaps conferring a degree of resilience to the antigenic shifts of SARS-CoV-2.

### Limitations of the study

In the present study, we report two broadly neutralizing antibodies that target a novel quaternary NTD-SD1 epitope. Although 12-16 and 12-19 showed robust activity against SARS-CoV-2 in vivo in hamster challenge experiments, the use of these mAbs as treatment was not examined. Additionally, the Fc-mediated effector functions of these two mAbs have not been evaluated.

## ACKNOWLEDGMENTS

This study was financially supported by the SARS-CoV-2 Assessment of Viral Evolution Program, NIAID, NIH (Subcontract No. 0258-A709-4609 under Federal Contract No. 75N93021C00014), the Gates Foundation (Project No. INV019355), Health@InnoHK, Innovation and Technology Commission, the Government of the Hong Kong Special Administrative Region, A. and P. Cherng, S. Yin, C. Ludwig, D. and R. Wu, and Regeneron Pharmaceuticals. We acknowledge D.S. Perlin for providing COVID-19 patient samples. Additionally, we thank C. Lu and the Columbia Center for Translational Immunology (CCTI) Flow Cytometry Core for cell sorting, the Columbia University Single Cell Analysis Core for 10X Genomics sequencing, and B. Grassucci and Z. Zhang for cryo-EM data collection assistance at the Columbia University Cryo-EM Facility. B.D. and J.D.B. were supported in part by NIH/NIAID grant R01AI141707, and J.D.B. is an Investigator of the Howard Hughes Medical Institute. We also appreciate the data contributors who shared their genomic data via GISAID.

## AUTHOR CONTRIBUTIONS

D.D.H. and L.L. conceived this project. L.L. and S.I. sorted antigen-specific memory B cells. Y.G. and Z.S. analyzed the 10X Genomics sequencing data and antibody repertoire. J.Y. cloned and expressed antibodies. L.L. expressed and purified spike proteins. L.L. conducted ELISA binding, SPR, cell surface staining, and competition experiments. L.L., Q.W., and S.I. performed pseudovirus neutralization experiments. M.S.N., Y.H., and H.M conducted authentic SARS-CoV-2 neutralization experiments. R.G.C. solved the cryo-EM structures. Y.G., R.G.C., E.R.R., and L.S. conducted the structural analyses. L.L., Q.W., and Y.G. design and conduct the S1 shedding experiment. J.F.-W.C., S.Y., V.K.-M.P., C.C.-S.C., and K.-Y.Y. conducted the SARS-CoV-2 in vivo experiments. B.D. and J.D.B. performed and analyzed the pseudovirus-based mutational scanning assay. L.L., Y.G., R.G.C., Q.W., S.I., B.D., L.S., and D.D.H. analyzed the results and wrote the manuscript. All authors approved the manuscript.

## SUPPLEMENTARY TABLE AND FIGURE LEGENDS

Table S1. Subject information. See also Figure 1.

Table S2. CryoEM Data Collection and Model Refinement. See also Figure 2.

Movie 1. Cryo-EM 3D variability analysis of the 12-19 3-Fab-bound spike particle class. See also Figure 3.

Movie 2. Cryo-EM 3D variability analysis of the 12-19 2-Fab-bound spike particle class. See also Figure 3.

Figure S1. Neutralization potency of Patient 12 serum and the isolated mAbs.

**(A)** Neutralization curves and ID_50_ titers of the convalescent serum from Patient 12 against pseudotyped SARS-CoV-2 variants and SARS-CoV.

**(B)** Neutralization curves of 27 isolated mAbs from Patient 12 against D614G and BA.1 pseudotyped viruses.

**(C)** Neutralization IC_50_ titers summarized from panel **B**.

**(D)** Neutralization curves of 12-16, 12-19, and S309 against authentic SARS-CoV-2 WA1 and BA.1.

**(E)** Neutralization curves of 12-16, 12-19, and S309 against a panel of SARS-CoV-2 pseudoviruses.

Data are representative of those obtained in three independent experiments and data and shown as mean ± SEM.

See also **Figure 1**.

Figure S2. Genetic analysis for mAbs12-16 and 12-19.

**(A)** Germline gene assignment for mAbs 12-16 and 12-19.

**(B)** Gene-specific substitution profile (GSSP) for mAb 12-16.

**(C)** Gene-specific substitution profile (GSSP) for mAb 12-19. For (B) and (C), the dots represent the conserved residues in antibody sequence, as compared with the germline gene. The CDRs are highlighted by rectangles.

**(D)** CDRH3 VDJ junction analysis for mAbs 12-16 and 12-19. Germline nucleotides and amino acid residues are shown in black with the corresponding junctions colored in light blue. Somatic hypermutations are colored in red. Nucleotides deleted by exonuclease trimming are indicated with strikethrough. The blue nucleotides represent the N and P nucleotide additions at the junctions.

See also **Figure 1**.

Figure S3. Cryo-EM data for mAbs 12-16 (A - E) or 12-19 (F - K) in complex with SARS-CoV-2 D614G spike trimer.

**(A)** and **(F)** Micrograph power spectrum (left) with contrast transfer function fit (right).

**(B)** and **(G)** Global refinement Fourier Shell Correlation curve showing overall resolution.

**(C)** and **(H)** Map density shown as mesh for mAb CDRH3 loop showing side chain fits.

**(D)** and **(I)** Local resolution mapped onto global refinement reconstruction.

**(E)** and **(K)** Local resolution for the antibody interfaces, shown from top and bottom. See also **Figure 2**.

**Figure S4. Structural comparison of mAbs 12-16 and 12-19 bound to SD1 and NTD.** The difference in angle of approach between heavy chains of 12-16 and 12-19 is shown. The close-up view highlights the CDR loop contacts between the antibody light chains and the N4-loop. See also **Figure 2**.

Figure S5. Cryo-EM processing and particle classification pipeline.

**(A)** For 12-16, nearly all spike particles were classified as 3-Fab-bound.

**(B)** For 12-19, spike particles were classified as 3- or 2-Fab-bound. See also **Figure 3**.

Figure S6. 12-16 and 12-19 inhibit CB6-spike binding and CB6-induced S1 shedding.

**(A)** Antibodies in complex with SARS-CoV-2 spike with one RBD in the up position (PDB: 7KRR). All of these antibodies can neutralize SARS-CoV-2 except 4-33. CB6 targets the RBD in the “up” position exclusively.

**(B)** Competition assay of CB6 binding to cell surface-expressed SARS-CoV-2 D614G spike in the presence of competitor antibodies. The data are shown as the mean ± SEM.

**(C)** CB6-induced S1 shedding from SARS-CoV-2 virions. D614G pseudovirus particles were incubated with CB6 at different doses for one hour at 37°C before the retained S1 and S2 subunits were determined by western blot.

**(D)** Inhibition of CB6-induced S1 shedding from the spike trimers on SARS-CoV-2 virions by the indicated antibodies. D614G pseudovirus particles were incubated with the indicated antibodies for one hour prior to incubating with 1 µg/mL CB6 for another one hour. The retained S1 and S2 subunits were determined by western blot.

**(E)** The intensities of the S1 and S2 glycoprotein bands in **(D)** were measured and the S1/S2 ratios are shown. Numbers denote the concentration of each antibody which inhibited half of the shedding of S1.

The results in **B**, **C, D, and E** are representative of those obtained in two independent experiments. See also **Figure 4**.

Figure S7. Neutralization mechanism of 12-16 and 12-19.

**(A)** Structural modeling shows Fc and Fab of NTD supersite antibody clash with dimeric hACE2. The IgG structure was made by superimposing heavy chains of crystal IgG structure (PDB:1HZH) with supersite antibody (PDB: 7L2F).

**(B)** SDS-PAGE of purified IgG, F(ab’)2, and Fab.

**(C)** Binding of antibodies to SARS-CoV-2 D614G spike protein was determined by surface plasmon resonance (SPR). The spike protein-bound sensors were incubated with six different concentrations of antibodies. Kinetic data from one representative experiment were fit to a 1:1 binding model. Data are representative of one from two independent experiments.

See also **Figure 5**.

Figure S8. Key spike escape mutations of 12-19 identified by deep mutational scanning.

**(A)** Mean escape scores for 12-19 antibody at each site in the BA.1 spike.

**(B)** Surface representation of spike colored by mean of escape scores at that site. PDB ID: 6XR8. Site numbering is based on the Wuhan-Hu-1 sequence.

**(C)** Antibodies 12-16 and 12-19 in complex with NTD and SD1. The NTD antigenic supersite is outlined in cyan. The mutations observed in VOCs are denoted as red spheres. Omicron BA.1, BA.2 (or BA.4/5, BQ.1), XBB.1, and shared mutations in at least two variants are colored in red, blue, green, and magenta, respectively.

See also **Figure 6**.

## STAR METHODS

### KEY RESOURCE TABLE

#### RESOURCE AVAILABILITY

##### Lead contact

Further information and requests for resources and reagents should be directed to and will be fulfilled by the lead contact, Dr. David D. Ho.

##### Materials availability

All requests for resources and reagents should be directed to and will be fulfilled by the Lead Contact, Dr. David D. Ho. This includes selective cell lines, plasmids, antibodies, viruses, sera, and proteins. All reagents will be made available on request after completion of a Material Transfer Agreement.

##### Data and code availability

- Paired heavy and light chain sequences for 12-16, 12-19, and 4-33 have been deposited to GenBank under accession XXXXXX to XXXXXX. The cryo-EM structures and the crystallographic structure are in the process of being deposited to the Electron Microscopy Data Bank (EMDB) and the Protein Data Bank (RCSB PDB). Cryo-EM structural models and maps of NTD-SD1- and NTD-directed antibodies in complex with SARS-CoV-2 spike have been deposited in the PDB and EMDB for antibodies 12-16 (PDB: 7UKL, EMDB: 26583), 12-19 (PDB: 7UKM, EMDB:26584), 4-33 (PDB: 8CSJ, EMDB: 26964). A summary of model refinement statistics is shown in **Table S2**.
- The computer code used to analyze the deep mutational scanning data is available at https://github.com/dms-vep/SARS-CoV-2_Omicron_BA.1_spike_DMS_12-19 and interactive display of the results is available at https://dms-vep.github.io/SARS-CoV-2_Omicron_BA.1_spike_DMS_12-19/.
- Any additional information required to reanalyze the data reported in this paper is available from the lead contact upon request.

### EXPERIMENTAL MODEL AND SUBJECTS

#### Human subjects

The protocol for sample acquisition was reviewed and approved by the Hackensack Meridian School of Medicine (Protocol No. Pro2020-0633). Informed consent was obtained from Patient 12, who was symptomatic for COVID-19 in November 2020. Following this, the patient received two doses of the mRNA-1273 vaccine in January and February 2021, and a blood sample was collected one week after the second vaccination. Sequencing analysis confirmed that Patient 12 was infected with the R.1 variant (B.1.1.316.1) of SARS-CoV-2.

#### Cell lines

Vero-E6 cells (CRL-1586) and HEK293T cells (CRL-3216) were obtained from the ATCC, while Expi293 cells (A14527) were obtained from Thermo Fisher Scientific. The morphology of each cell line was visually confirmed prior to use, and all cell lines tested negative for mycoplasma contamination. Vero-E6 cells originated from African green monkey kidneys, while HEK293T and Expi293 cells were of female origin.

### METHOD DETAILS

#### Plasmid constructs

The constructs used for expression of SARS-CoV-2 variant spikes and stabilized soluble SARS-CoV-2 S2P spike trimer proteins were obtained from previous studies^9–13, 47, 62^. For antibody expression, the variable genes of antibodies were optimized for eukaryotic cell expression and synthesized by GenScript. Genes for the variable regions were then separately inserted into expression vectors (gWiz or pcDNA3.4) containing the corresponding constant region for heavy and light chains.

#### Expression and purification of SARS-CoV-2 S2P spike trimer proteins

Expi293 cells were used for transient transfection with the suitable S2P stabilized spike-expression vector by using 1 mg/mL polyethylenimine (PEI, Polysciences). The supernatant was harvested and the spike trimer was purified using Ni-NTA resin (Invitrogen) in accordance with the manufacturer’s protocol five days after transfection. Prior to use, all proteins were evaluated for size and purity via SDS-PAGE.

#### Sorting for S trimer-specific B cells and single-cell B cell receptor sequencing

S2P spike trimer-specific memory B cells were isolated and sequenced using the protocol previously described by Liu et al.^21, 27^. In brief, peripheral blood mononuclear cells (PBMCs) from patient 12 were stained with LIVE/DEAD Fixable Yellow Dead Cell Stain Kit (Invitrogen) at room temperature for 20 min, washed with RPMI-1640 complete medium, and incubated with 10 μg/mL B.1.351 S2P spike trimer at 4_°C for 45 min. Cells were then washed and incubated with a cocktail of flow cytometry and hashtag antibodies, containing CD3 PE-CF594 (BD Biosciences), CD19 PE-Cy7 (Biolegend), CD20 APC-Cy7 (Biolegend), IgM V450 (BD Biosciences), CD27 PerCP-Cy5.5 (BD Biosciences), anti-His PE (Biolegend), and human Hashtag 3 (Biolegend) at 4_°C for 1 h. Cells were then washed again, resuspended in RPMI-1640 complete medium, and sorted for S2P spike trimer-specific memory B cells (CD3−CD19+CD27+S trimer+ live single lymphocytes). These sorted cells were mixed with PBMCs from the same donor, labeled with Hashtag 1, and loaded to a 10X Chromium chip for the 5′ Single Cell Immune Profiling Assay (10X Genomics) at the Columbia University Human Immune Monitoring Core (HIMC; RRID:SCR_016740). Library prep and quality control were performed according to the manufacturer’s instructions and then sequenced on a NextSeq 500 (Illumina).

#### Identification of S trimer-specific antibody transcripts

S2P spike trimer-specific antibody transcripts were identified as previously described^21, 27^. Full-length antibody transcripts were assembled using the Cell Ranger V(D)J pipeline (version 3.1.0, 10X Genomics) with default parameters using the GRCh38 V(D)J germline sequence version 2.0.0 as the reference. To distinguish cells from the antigen sort and spike-in, we first used the count module in Cell Ranger to calculate copies of all hashtags in each cell from the NGS raw reads. High-confidence antigen-specific cells were identified as follows: 1. A cell must contain more than 100 copies of the antigen sort-specific hashtag to qualify as an antigen-specific cell. 2. As hashtags can fall off cells and bind to cells from a different population in the sample mixture, each cell usually has both sorted and spiked-in-specific hashtags. To enrich for true antigen-specific cells, the copy number of the specific hashtag is set such that it has to be at least 1.5× higher than that of the non-specific hashtag. 3. Low-quality cells were identified and removed using the cell-calling algorithm in Cell Ranger. 4. Cells that did not have productive heavy and light chain pairs were excluded. 5. If a cell contained more than two heavy and/or light chain transcripts, the transcripts with fewer than three unique molecular identifiers were removed. 6. Cells with identical heavy and light chain sequences, which may be from mRNA contamination, were merged into one cell.

#### Antibody transcript annotation

The transcripts of antigen-specific antibodies were subjected to quality control and annotation using SONAR version 2.0, as previously described^21, 27, 63^. The V(D)J genes were assigned to each transcript using BLASTn with customized parameters against a germline gene database obtained from the international ImMunoGeneTics information system (IMGT) database^64^. The CDR3 was identified based on the BLAST alignments of the V and J regions, using the conserved second cysteine in the V region and WGXG (heavy chain) or FGXG (light chain) motifs in the J region, where X represents any amino acid. To assign the isotype for heavy chain transcripts, the constant domain 1 (CH1) sequences were used, and a database of human CH1 genes obtained from IMGT was searched using BLASTn with default parameters. The CH1 allele with the lowest E-value was used for significant isotype assignments, using a BLAST E-value threshold of 10-6. Transcripts containing incomplete V(D)J and/or frameshifts were excluded, and sequences other than the V(D)J region were removed. The remaining transcripts were aligned to the assigned germline V gene using CLUSTALO, and the somatic hypermutation level was calculated using the Sievers method^65^. The D gene assignment for each transcript was performed using the default parameters of the HighV-QUEST function in the IMGT web server. In cases where cells had multiple high-quality heavy or light chains, which may have arisen from doublets, all H and L chain combinations were synthesized.

#### Antibody expression and purification

Expi293 cells were utilized to co-transfect the antibody heavy chain and light chain expressing plasmids. Transfection was facilitated using 1 mg/mL polyethylenimine (PEI) and the cells were cultured with shaking at 125 rpm under 8% CO2 at 37_°C. After five days of transfection, the antibody was purified from the supernatant using rProtein A Sepharose, following the instructions provided by the manufacturer.

#### F(ab’)2 and Fab preparation

To generate F(ab’)2 format antibodies, the full-length IgG was digested using immobilized pepsin protease with the Pierce™ F(ab’)2 Preparation Kit (ThermoFisher, Cat# 44988) according to the manufacturer’s instructions. The production of Fab fragments from IgG antibodies was achieved through the utilization of immobilized Endoproteinase Lys-C (Sigma-Aldrich). The digestion was performed in a buffer consisting of 25 mM Tris (pH 8.5) and 1 mM EDTA for a duration of 3 hours. The Fab fragments were purified from the cleaved Fc domain through affinity chromatography using rProtein A Sepharose. The purity of the IgG, F(ab’)2, and Fab antibodies was assessed by sodium dodecyl sulfate-polyacrylamide gel electrophoresis (SDS-PAGE) using a Bis-Tris protein gel (Invitrogen) and a MES buffer system, prior to any experimental use.

#### Epitope and paratope analysis

The paratope and epitope residues for 12-16 and 12-19 were identified using PISA with the default parameters^66^. The combined Fab interface model (Fab+SD1+RBD+NTD) was saved and imported into PyMOL and the Interface residues script was run using a dASA cutoff of 0.75 Å2. The gene-specific substitution profiles (GSSP) for 12-16 and 12-19 germline genes were obtained from the cAb-Rep database (https://cab-rep.c2b2.columbia.edu/)^67^.

#### C**α** distance calculation

The up and down spike protomers were first superimposed by using the ‘align’ function in PyMOL 2.3.2. The distance between the Cαs from identical residues within the two protomers was then determined using the rms_cur function in PyMOL.

#### Production of pseudoviruses

The generation of recombinant Indiana vesicular stomatitis virus (rVSV) pseudotyped^21^. Briefly, HEK 293T cells were transfected with the relevant SARS-CoV-2 spike expression construct using 1 mg/mL polyethyleneimine (PEI). The following day, the cells were infected with VSV-G pseudotyped ΔG-luciferase (G*ΔG-luciferase, Kerafast) at a multiplicity of infection (MOI) of 3 for 2 hours. The cells were then washed three times with fresh medium and incubated for an additional 24 hours. The supernatant was collected, cleared through centrifugation, and aliquoted and stored at −80°C until usage. The titer of all pseudoviruses was determined to normalize the viral input prior to use in subsequent assays.

##### Pseudovirus neutralization assay

Pseudovirus neutralization assays were conducted by incubating pseudoviruses with serial dilutions of convalescent sera from COVID-19 patients, human ACE2-Fc, or antibodies (IgG, F(ab’)2, or Fab) in triplicate in 96-well plates. The incubation was performed at 37°C for 1 hour. Subsequently, 3 x 10^4^ Vero-E6 cells were added to each well and incubated for an additional 10-12 hours. The cells were then lysed, and luciferase activity was quantified using the Luciferase Assay System (Promega) following the manufacturer’s protocol. The neutralization activity was calculated based on the reduction in luciferase activity compared to the mock-treated control wells. The concentrations of mAbs and dilutions of serum that inhibited 50% of infection (IC_50_ and ID_50_ values, respectively) were determined by fitting the data to five-parameter dose-response curves using GraphPad Prism 9 (GraphPad Software Inc.).

#### Neutralization of authentic SARS-CoV-2 by purified mAbs

The neutralizing activity of authentic SARS-CoV-2 was determined through an endpoint dilution assay performed in a 96-well plate format, as previously described^21, 27^. mAbs were serially diluted and incubated with SARS-CoV-2 at a multiplicity of infection (MOI) of 0.1 in Eagle’s minimum essential medium (ATCC) supplemented with 7.5% inactivated FBS for 1 hour at 37°C in triplicate. The antibody-virus mixture was then added to a monolayer of Vero-E6 cells that had been grown overnight, and the cells were incubated for 70 hours. The morphological changes resulting from cytopathic effects (CPE) due to virus infection were visually scored for each well on a scale of 0 to 4, with 4 indicating complete virus-mediated cytopathy. The scoring was performed in a double-blinded manner, and the results were converted into a percentage of neutralization. The IC_50_ was determined by fitting the data to five-parameter dose-response curves using GraphPad Prism 9.

#### Antigen binding testing by ELISA

Binding of antibodies to S trimer was tested in enzyme-linked immunosorbent assay (ELISA) as previously described^21, 27^. Briefly, 50 ng of S trimer per well was coated onto ELISA plates at 4 °C overnight. Plates were washed, then blocked with 300 μL of blocking buffer (PBS + 1% bovine serum albumin + 20% bovine calf serum (Sigma-Aldrich)) at 37°C for 2 h. Plates were again washed, then 100 μL of the serially diluted antibodies were added and incubated at 37°C for 1 h, washed again, and then 100 μL of Peroxidase AffiniPure goat anti-human IgG Fcγ fragment–specific antibody (Jackson ImmunoResearch, catalog no. 109-035-170, RRID: AB_2810887, 1:10,000 dilution in blocking buffer) was added and incubated for 1 h at 37°C. Finally, 3,3′,5,5′- tetramethylbenzidine (TMB) substrate (Sigma-Aldrich) was added to initiate the reaction and stopped using 1 M sulfuric acid. Absorbance for all wells was then measured at 450 nm. The concentration of antibody that gives half-maximal binding (EC50) was determined by fitting the data to five-parameter dose-response curves in GraphPad Prism 9.

#### Cell-surface S protein binding and competition binding assays

Binding of antibodies S trimers expressed on cell surfaces was tested as previously described^21, 27^. Expi293 cells were first co-transfected with pRRL-cPPT-PGK-GFP (Addgene) and the appropriate SARS-CoV-2 spike expression vector at a ratio of 1:1 using 1 mg/mL PEI and incubated at 37 °C shaking at 125 rpm under 8% CO2 for 48 h. Cells were then harvested and incubated with 12-16, 12-19, or S309 at a final concentration of 10 µg/mL at 4 °C for 45 min. Then, 100 μL of APC anti-human IgG Fc (BioLegend, catalog no. 366906, RRID: AB_2888847, 1:20 dilution) was added and incubated at 4°C for 45 min. Cells were washed three times with FACS buffer (PBS + 2% FBS) before each step. Cells were then resuspended, and antibody binding was quantified by flow cytometry on a LSRII (BD Biosciences). The mean fluorescence intensity of antibody-bound APC-positive cells within green fluorescent protein (GFP)- positive cells was determined using FlowJo (BD Biosciences).

For the ACE2 and CB6 competition binding assays, human ACE2-Fc (SinoBiological, Cat# 10108-H02H) and CB6 were biotinylated by One-Step Antibody Biotinylation Kit (Miltenyi Biotec, Cat# 130-093-385) according to the manufacturer’s instructions. The transfected Expi293 cells were then incubated with a mixture of biotinylated ACE2-Fc or CB6 (0.25 μg/mL) and serially diluted competitor antibodies at 4°C for 1 h. Afterwards, 100 μL of diluted APC-streptavidin (Biolegend) was added to the cells and incubated at 4°C for 45 min. Cells were washed three times with FACS buffer before each step. Finally, cells were resuspended and binding of ACE2-Fc or CB6 to cell-surface S trimer was quantified by flow cytometry on a LSRII (BD Biosciences). The mean fluorescence intensity of APC in GFP-positive cells was determined using FlowJo and the relative binding of ACE2-Fc or CB6 to the S trimer in the presence of competitors was calculated as the percentage of the mean fluorescence intensity compared to that of the competitor-free controls.

#### Hamster protection experiment

In vivo evaluation of mAbs 12-16 and 12-19 was performed in an established Syrian hamster model for COVID-19 as previously described with slight modifications^68^. At 24 h before SARS-CoV-2 Omicron (B.1.1.529.1 or BA.1) variant challenge, each hamster (n = 6 per group) was intraperitoneally administered with one dose of 10 mg/kg of 12-16, 12-19, or control HIV-1 neutralizing antibody (3BNC117) in phosphate-buffered saline (PBS). Twenty-four hours later, each hamster was intranasally inoculated with a challenge dose of 100 µL of Dulbecco’s Modified Eagle Medium containing 105 plaque-forming units of SARS-CoV-2 Omicron (hCoV-19/Hong_Kong/HKU-211129-001/2021; GISAID accession number: EPI_ISL_6841980) under intraperitoneal ketamine (200 mg/kg) and xylazine (10 mg/kg) anesthesia^69^. The hamsters were monitored daily for clinical signs of disease and sacrificed at 4 days post-challenge. Half of each hamster’s lung tissues were used for viral load determination by the quantitative COVID-19- RdRp/Hel reverse transcription-polymerase chain reaction assay^70^ and viral titer determination by plaque assay as previously described^71^. Unpaired t test was used to determine significant differences among the different groups. P values less than 0.05 were considered statistically significant.

#### Cryo-EM grid preparation

Samples for cryo-EM were prepared in a buffer of 10 mM sodium acetate, 150 mM NaCl, and adjusted to pH 5.5. Spike + Fab complex was made by mixing purified SARS-CoV-2 S2P D614G spike trimer protein with Fabs in a 1:3 molar ratio (spike protomer: Fab), such that the final concentration of spike was 1 mg/mL. This mixture was incubated on ice for 1 h. Before freezing, 0.005% (w/v) n-Dodecyl β-D-maltoside (DDM) was added to deter preferred orientation and aggregation during vitrification. Cryo-EM grids were prepared by applying 3 µL of sample to a freshly glow-discharged carbon-coated copper grid (CF 1.2/1.3 300 mesh); the sample was vitrified in liquid ethane using a Vitrobot Mark IV with a wait time of 30 s, a blot time of 3 s, and a blot force of 0.

#### Cryo-EM data collection and analysis

Cryo-EM data for single particle analysis were collected at the Columbia Cryo-EM Facility (12-16 and 12-19) and the Columbia Zuckerman Institute (4-33) on a Titan Krios electron microscope operating at 300 kV, equipped with a Gatan K3-BioQuantum direct detection detector and energy filter, using the Leginon^72^ (12-16 and 12-19) and SerialEM^73^ (4-33) software packages. For 12-16 and 12-19, exposures were taken at a magnification of 105,000x (pixel size of 0.83 Å), using a total electron fluence of 58 e-/Å2 fractionated over 50 frames with an exposure time of 2.5 s. For 4-33, exposures were taken at a magnification of 81,000x (pixel size of 1.07 Å), using a total electron fluence of 42 e-/Å2 fractionated over 60 frames with an exposure time of 3 s. A random defocus range of −0.8 to −2.0 µm was used.

Data processing was performed using cryoSPARC v3.3.1^74^. Raw movies were aligned and dose-weighted using patch motion correction, and the micrograph contrast transfer function (CTF) parameters were estimated using patch CTF estimation. Micrographs were picked using a blob picker and an initial spike particle set was selected using 2D classification. Ab-initio jobs were run using 1 class, as well as multiple classes to examine the diversity of spike conformations. Heterogenous refinement was used to remove junk particles and sort between 2-Fab-bound vs 3-Fab-bound classes, as well as RBD-up/down in the 2-Fab-bound particle set. Cryo-EM classification and processing details are shown in Figure S5. The resulting curated particle sets were local motion corrected and refined to high resolution using homogenous refinement. An additional boost in resolution was obtained by utilizing the “minimize over per-particle scale” parameter (starting at iteration 3), combined with the “optimize per particle defocus” and “optimize per-group CTF params” parameters in homogenous refinement. The default cryoSPARC auto-sharpened maps were used to build the models. Cryo-EM data collection and consensus refinements are summarized in Figure S3. Data collection and processing statistics are shown in **Table S2**.

#### 3D variability analysis

3D variability analysis was performed according to the cryoSPARC guide, which is described generally by (https://guide.cryosparc.com/processing-data/tutorials-and-case-studies/tutorial-3d-variability-analysis-part-one). Briefly, a number of 3D Variability jobs were run using either 3 or 4 modes and filter resolutions in the range of 5 Å to 10 Å. The RBD motion was captured well using 8 Å filter resolution. Results were visualized using the 3D Variability Display job run in simple mode using 5 frames, with filter resolutions between 4 Å and 8 Å. The movies were colored by fitting each colored domain into the 5 frames of the movie (each frame is an intermediate reconstruction) using Chimera’s^75^ “fit to map” tool followed by the “scolor” command. Movies were generated using the volume series feature using the script provided by cryoSPARC. Playback speed was slowed down by including “framerate 5” parameter in the “movie encode” command. Top and side view movies were concatenated, and labels were added using Windows Movie Maker.

#### Model building and refinement

Initial molecular models for Fabs were generated using Alphafold multimer^76^ using paired Heavy and Light sequences. A 3-RBD-down spike model from (pdb: 6XM5) was used as a starting model. The initial models were rigid body docked into the density map using Chimera’s “fit to map” tool and combined. The Fab CDR loops were manually fit into the density map using Coot^77^. The models were fit into density using the ISOLDE^78^ package in ChimeraX^79^. Ramachandran outliers were corrected using ISOLDE’s “flip peptide bond” feature. The resulting models were then refined using Rosetta’s “Relax” script^80–83^. Finally, real space refinement in Phenix^84^ was performed to remove geometry outliers. The remaining manual adjustments were performed in Coot. Models were validated using Molprobity^85^ in Phenix and the PDB validation server, and deposited to the PDB with accession codes: 7UKL (12-16), 7UKM (12-19), 8CSJ (4-33). Maps were deposited to the EMDB with codes: 26583 (12-16), 26584 (12-19), 26964 (4- 33). A summary of model refinement statistics is shown in **Table S2**.

#### S1 shedding from spike trimers

D614G S glycoprotein pseudotyped VSV particles were first generated as described above and the evaluation of S1 subunit shedding was performed as previously described^54, 55^. To evaluate the induction of S1 shedding by hACE2-Fc (SinoBiological, Catalog #10108-H02H) and CB6, the cell supernatants containing pseudovirus particles were filtered through a 0.45 µm filter and then incubated with ACE2-Fc or CB6 at the appropriate concentrations at 37°C for 1 hour. The virus particles were then pelleted at 18,000 x g for 1 hour at 4°C. The pelleted virus particles were resuspended in 1 X LDS sample buffer (ThermoFisher, Cat# NP0008) and analyzed by Western blotting. The S1 and S2 subunits were detected using rabbit anti-SARS-Spike S1 (Sino Biological, Cat #40591-T62) and rabbit anti-SARS-Spike S2 (Sino Biological, Cat #40590-T62) primary antibodies, respectively. The Western blots were developed using horseradish peroxidase-conjugated anti-rabbit antibody (Cytiva, Cat #NA934-1ML). To test the inhibition of hACE2-Fc or CB6-induced S1 shedding from the spike trimer on SARS-CoV-2 virions by antibodies, SARS-CoV-2 D614G pseudovirus was first pretreated with different doses of selected antibodies for 1 hour at 37°C, then incubated with 5 µg/mL ACE2-Fc or 1 µg/mL CB6 for an additional hour at 37°C. The virus particles were then pelleted and analyzed by Western blotting as described above. The band intensities of S1 and S2 from unsaturated Western blots were determined using ImageJ software^86^, and the resulting ratio was plotted.

#### Antigen binding testing by SPR

The binding of antibodies (IgG, F(ab’)2, and Fab) to the SARS-CoV-2 spike trimer protein was assessed using Surface Plasmon Resonance (SPR) as previously described^27^. The SPR binding assays were conducted using a Biacore T200 biosensor with a Series S CM5 chip (Cytiva) in a running buffer consisting of 10 mM HEPES (pH 7.4), 150 mM NaCl, 3 mM EDTA, and 0.05% P-20 (HBS-EP+ buffer, Cytiva) at 25°C. The SARS-CoV-2 D614G spike protein stabilized by S2P was captured on the biosensor surface using an anti-His antibody surface, which was generated using a His-capture kit (Cytiva) according to the manufacturer’s instructions. The resulting antibody surface contained approximately 10,000 resonance units (RUs) of anti-His antibody per surface. The spike protein was captured on a single flow cell at 125 to 200 RUs. An anti-His antibody surface was used as a reference flow cell to eliminate bulk shift changes from the binding signal. The antibodies were evaluated using a three-fold dilution series with concentrations ranging from 1.2 to 33.3 nM. The association and dissociation rates were monitored for 55 seconds and 300 seconds, respectively, at a flow rate of 50 mL/min. The bound spike protein-antibody complexes were regenerated from the anti-His antibody surface using 10 mM glycine (pH 1.5). A blank buffer cycle was performed by injecting running buffer instead of antibody to eliminate systematic noise from the binding signal. The data obtained were processed and fitted to a 1:1 binding model using Biacore Evaluation Software.

#### Lentivirus-based full spike deep mutational scanning

The BA.1 full spike deep mutational scanning libraries have been previously described^57^. These libraries have ∼7000 mostly functional amino-acid mutations across all of the spike protein. Antibody escape mapping experiments for 12-19 antibody were performed as described^57^. Different concentrations of 12-19 antibody were incubated with the library virus for 1 h at 37°C. Antibody concentrations used for selection experiments were determined by running a pseudovirus neutralization assay on ACE2 overexpressing HEK-293T cells and selecting a starting concentration close to IC99 (12 µg/ml) and then increasing this concentration by four and sixteen-fold. After virus-antibody incubation, HEK-293T cells expressing medium amounts of ACE2 as described^87^ were infected. 15 h after infection viral genomes were harvested from cells and barcodes were sequenced as described^57^. Two biological library replicates (using independent mutant libraries) were used to map escape mutations.

Escape for each mutation in the library was calculated relative to a non-neutralized VSV-G pseudotyped standard as described^57^. This analysis uses a biophysical model described previously^88^, which is implemented in polyclonal package (https://jbloomlab.github.io/polyclonal/). The full analysis pipeline for 12-19 antibody and the underlying data can be found at https://dms-vep.github.io/SARS-CoV-2_Omicron_BA.1_spike_DMS_12-19 and interactive escape plots for this antibody can be found at https://dms-vep.github.io/SARS-CoV-2_Omicron_BA.1_spike_DMS_12-19/mAb_12-19_escape_plot.html.

### QUANTIFICATION AND STATISTICAL ANALYSIS

The p-values in **Figure 1** were determined through an unpaired t test. The levels of significance are indicated as follows: *p<0.05; **p<0.01; ***p<0.001; and ****p<0.0001. The EC_50_, IC_50_, and ID_50_ values were determined by fitting the data to five-parameter dose-response curves using GraphPad Prism 9. The Western blot data were analyzed using ImageJ software. FACS analysis was performed using a LSRII flow cytometer, and the data were analyzed using FlowJo 10 software. SPR data were processed and fitted to a 1:1 binding model using Biacore Evaluation Software, and the results were plotted using GraphPad Prism 9. The data presented are representative or mean data derived from at least two independent experiments.

