## Supplemental figures for "Antibodies that neutralize all current SARS-CoV-2 variants of concern by conformational locking"

### Table S1

| Sample ID | Symptoms | Infection & vaccination history | Blood collection days post 2nd vaccination |
|---|---|---|---|
| Patient 12 | Shortness of breath, fatigue, muscle pain, joint pain, headache, lack of taste, lack of smell, fever, and confusion | R.1 (B.1.1.316.1)/ mRNA-1273/mRNA-1273 | 7 |

### Table S2

| | SARS-CoV-2 S2P + 12-16 Fab | SARS-CoV-2 S2P + 12-19 Fab | SARS-CoV-2 S2P + 4-33 Fab |
|---|---|---|---|
| **EMDB ID** | 26583 | 26584 | 26964 |
| **PDB ID** | 7UKL | 7UKM | 8CSJ |
| **Data Collection** | | | |
| Microscope | FEI Titan Krios | FEI Titan Krios | FEI Titan Krios |
| Voltage (keV) | 300 | 300 | 300 |
| Magnification | 105k | 105k | 81k |
| Defocus Range (µm) | -0.8/-2.0 | -0.8/-2.0 | -0.8/-2.0 |
| Camera | Gatan K3 BioQuantum | Gatan K3 BioQuantum | Gatan K3 BioQuantum |
| Pixel Size (Å/pix) | 0.83 | 0.83 | 1.07 |
| Recording Mode | counting | counting | counting |
| Dose Rate (e-/pixel/s) | 16 | 16 | 16 |
| Electron Dose (e-/Å$^2$) | 58 | 58 | 42 |
| **Data Processing** | | | |
| Software | cryoSPARC v3.3 | cryoSPARC v3.3 | cryoSPARC v3.3 |
| Micrographs used | 3,691 | 4,042 | 6,100 |
| Number of Particles | 463,850 | 256,243 | 420,343 |
| Symmetry | C1 | C1 | C3 |
| Box Size (pix) | 440 | 440 | 384 |
| Global Map FSC0.143 (Å) | 3.09 | 3.03 | 3.56 |
| **Refinement and Validation** | | | |
| Software | Phenix, ISOLDE, Rosetta | Phenix, ISOLDE, Rosetta | Phenix, ISOLDE |
| Initial Model Used | 6XM5(Spike), Alphafold(Fab) | 6XM5(Spike), Alphafold(Fab) | 7LAB(Spike), SAbPred(Fab) |
| Number of Atoms | 30,234 | 30,145 | 32,073 |
| Protein Residues | 3,795 | 3,771 | 4,038 |
| Ligands | NAG: 48 | NAG: 47 | NAG: 53 |
| Model vs. Data CC (mask) | 0.79 | 0.87 | 0.7 |
| RMS deviations | | | |
| Bond lengths (Å) (# >4σ) | 0 | 0 | 1 |
| Bond angles (°) (# >4σ) | 17 | 2 | 65 |
| MolProbity Score | 1.25 | 1.07 | 1.96 |
| Clashscore (all atom) | 2.42 | 1.41 | 3.57 |
| Poor rotamers (%) | 0 | 0 | 0 |
| Ramachandran plot | | | |
| Favored (%) | 96.55 | 96.84 | 92.95 |
| Allowed (%) | 3.45 | 3.16 | 7.05 |
| Outliers (%) | 0 | 0 | 0 |

### Figure S1

A

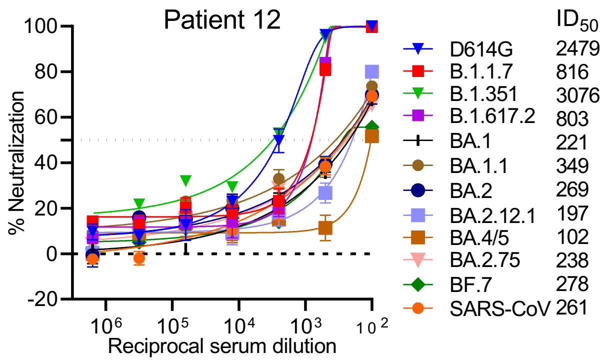


B

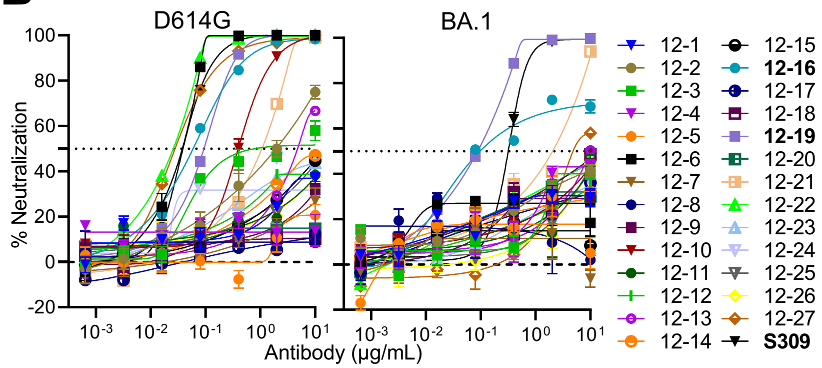


C

| IC50 (μg/mL) | D614G | BA.1 | IC50 (μg/mL) | D614G | BA.1 |
|---|---|---|---|---|---|
| 12-1 | >10 | >10 | 12-15 | >10 | >10 |
| 12-2 | 1.942 | >10 | 12-16 | 0.059 | 0.097 |
| 12-3 | 0.860 | >10 | 12-17 | >10 | >10 |
| 12-4 | >10 | >10 | 12-18 | >10 | >10 |
| 12-5 | >10 | >10 | 12-19 | 0.094 | 0.088 |
| 12-6 | 0.038 | >10 | 12-20 | >10 | >10 |
| 12-7 | >10 | >10 | 12-21 | 1.069 | 2.051 |
| 12-8 | >10 | >10 | 12-22 | 0.026 | >10 |
| 12-9 | >10 | >10 | 12-23 | >10 | >10 |
| 12-10 | 0.396 | >10 | 12-24 | >10 | >10 |
| 12-11 | >10 | >10 | 12-25 | >10 | >10 |
| 12-12 | >10 | >10 | 12-26 | >10 | >10 |
| 12-13 | 4.789 | 9.910 | 12-27 | 0.028 | 4.839 |
| 12-14 | >10 | >10 | S309 | 0.039 | 0.318 |

D

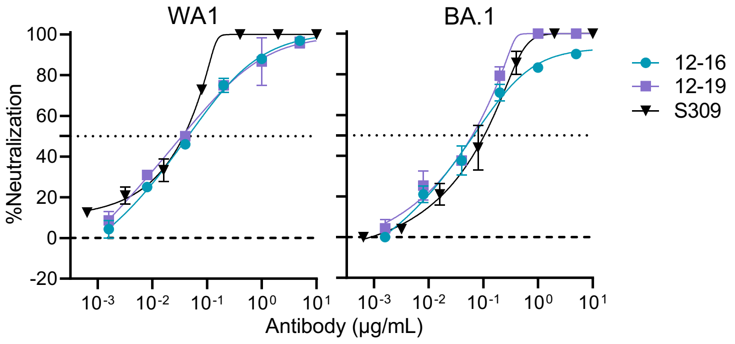


E

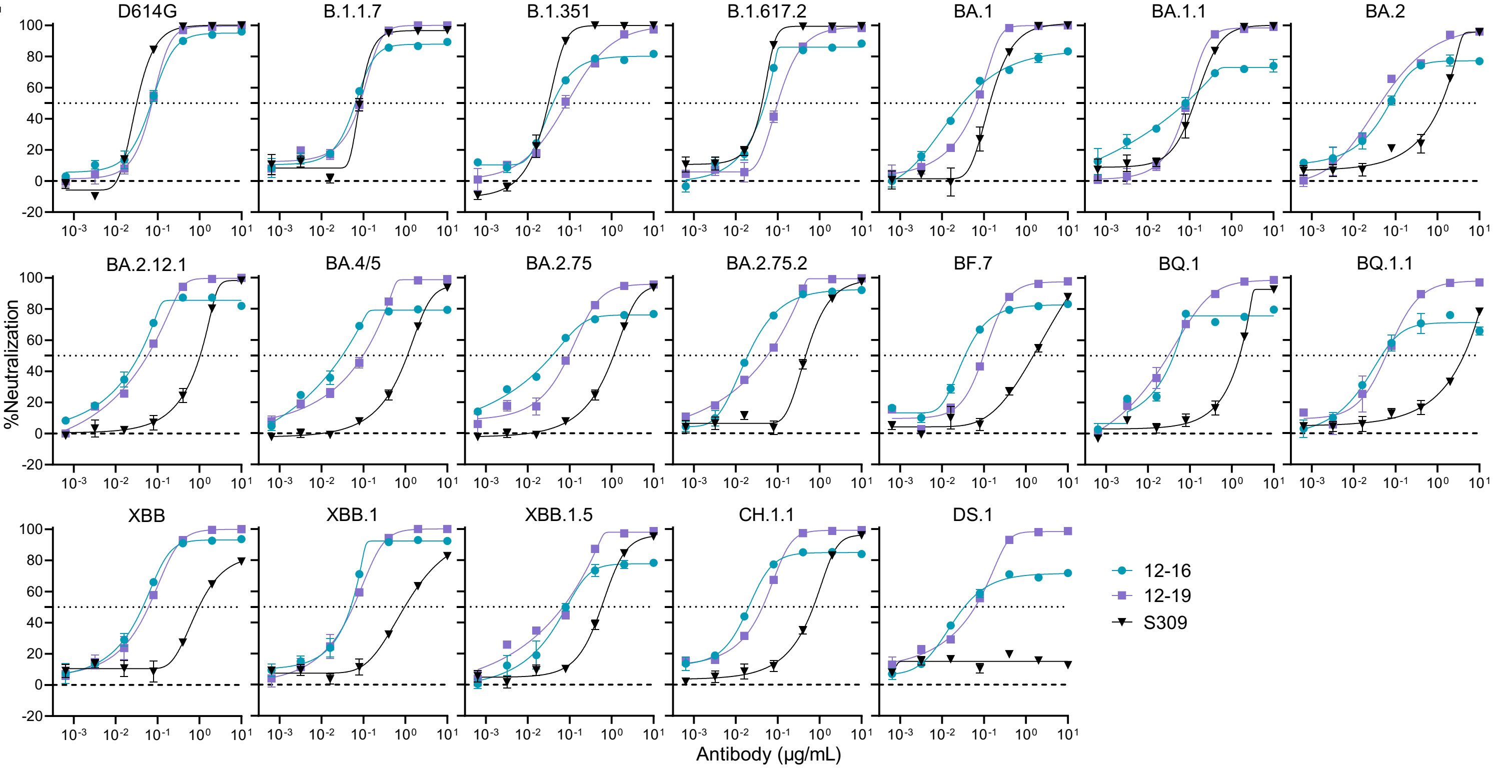

### Figure S2

**A**

| mAbs | HV | HD | HJ | Isotype | H_SHM | CDRH3_len | HCDR3 | LV | LJ | L_SHM | CDRL3_len | LCDR3 |
|---|---|---|---|---|---|---|---|---|---|---|---|---|
| 12-16 | IGHV3-30*18 | IGHD3-9*01 | IGHJ6*02 | IGHG3*04 | 1.70% | 25 | AKDLSFYYDISTGYYPQSYNSGTDV | IGLV3-1*01 | IGLJ*01 | 1.40% | 11 | QGWDRSTGYYV |
| 12-19 | IGHV3-33*01 | IGHD3-9*01 | IGHJ6*02 | IGHG1*04 | 2.70% | 26 | ARDRTPVYDILTGYYWPPPRVDGMDV | IGKV2-29*02 | IGKJ2*02 | 2.30% | 9 | MQSIQLPFT |

**B**

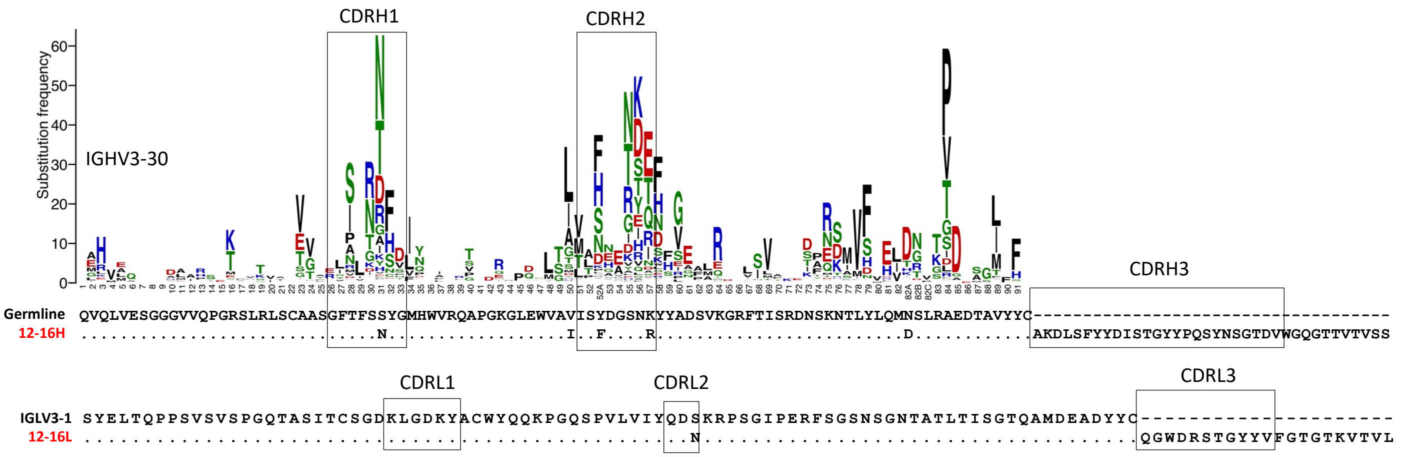


**C**

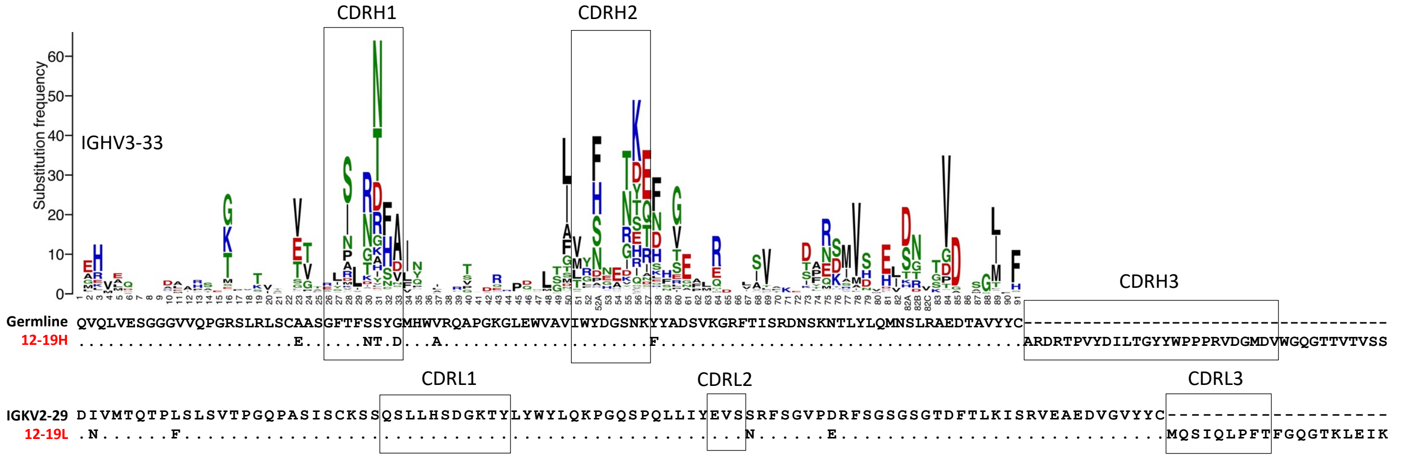


**D**

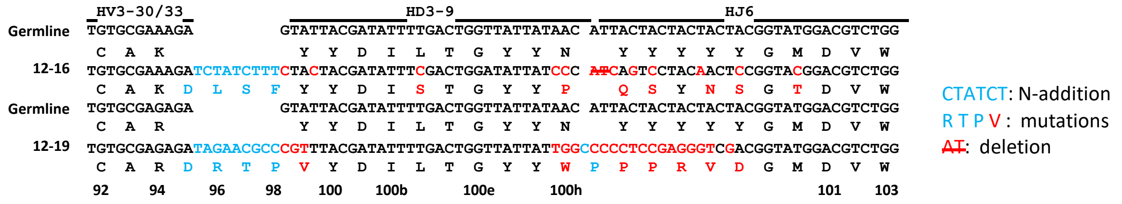

### Figure S3

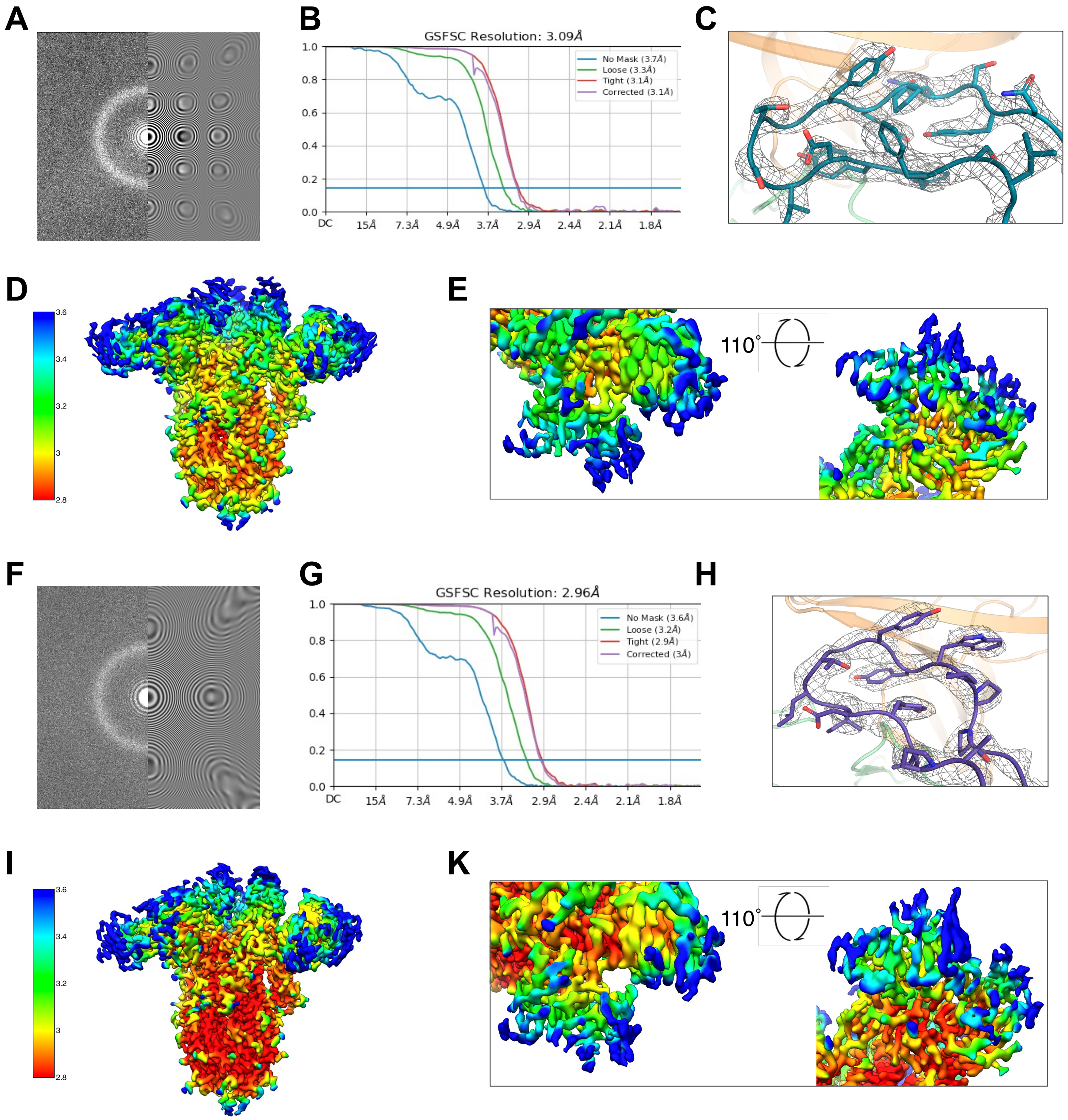

A
B
GSFSC Resolution: 3.09Å
No Mask (3.7Å)
Loose (3.3Å)
Tight (3.1Å)
Corrected (3.1Å)
1.0
0.8
0.6
0.4
0.2
0.0
DC
15Å
7.3Å
4.9Å
3.7Å
2.9Å
2.4Å
2.1Å
1.8Å
C
D
3.6
3.4
3.2
3
2.8
E
110°
F
G
GSFSC Resolution: 2.96Å
No Mask (3.6Å)
Loose (3.2Å)
Tight (2.9Å)
Corrected (3Å)
1.0
0.8
0.6
0.4
0.2
0.0
DC
15Å
7.3Å
4.9Å
3.7Å
2.9Å
2.4Å
2.1Å
1.8Å
H
I
3.6
3.4
3.2
3
2.8
K
110°

### Figure S4

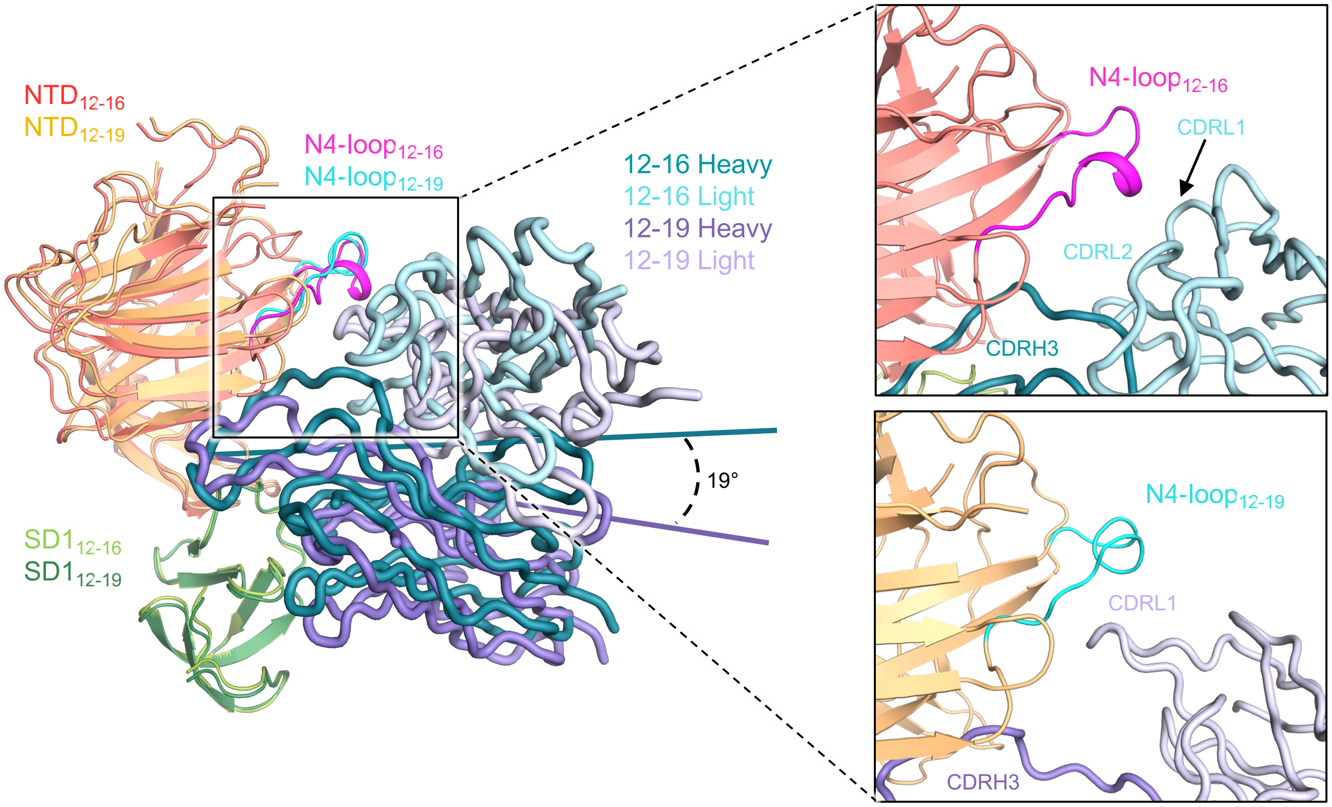

NTD12-16
NTD12-19
N4-loop12-16
N4-loop12-19
12-16 Heavy
12-16 Light
12-19 Heavy
12-19 Light
19°
SD1 12-16
SD1 12-19
N4-loop12-16
CDRL1
CDRL2
CDRH3
N4-loop12-19
CDRL1
CDRH3

### Figure S5

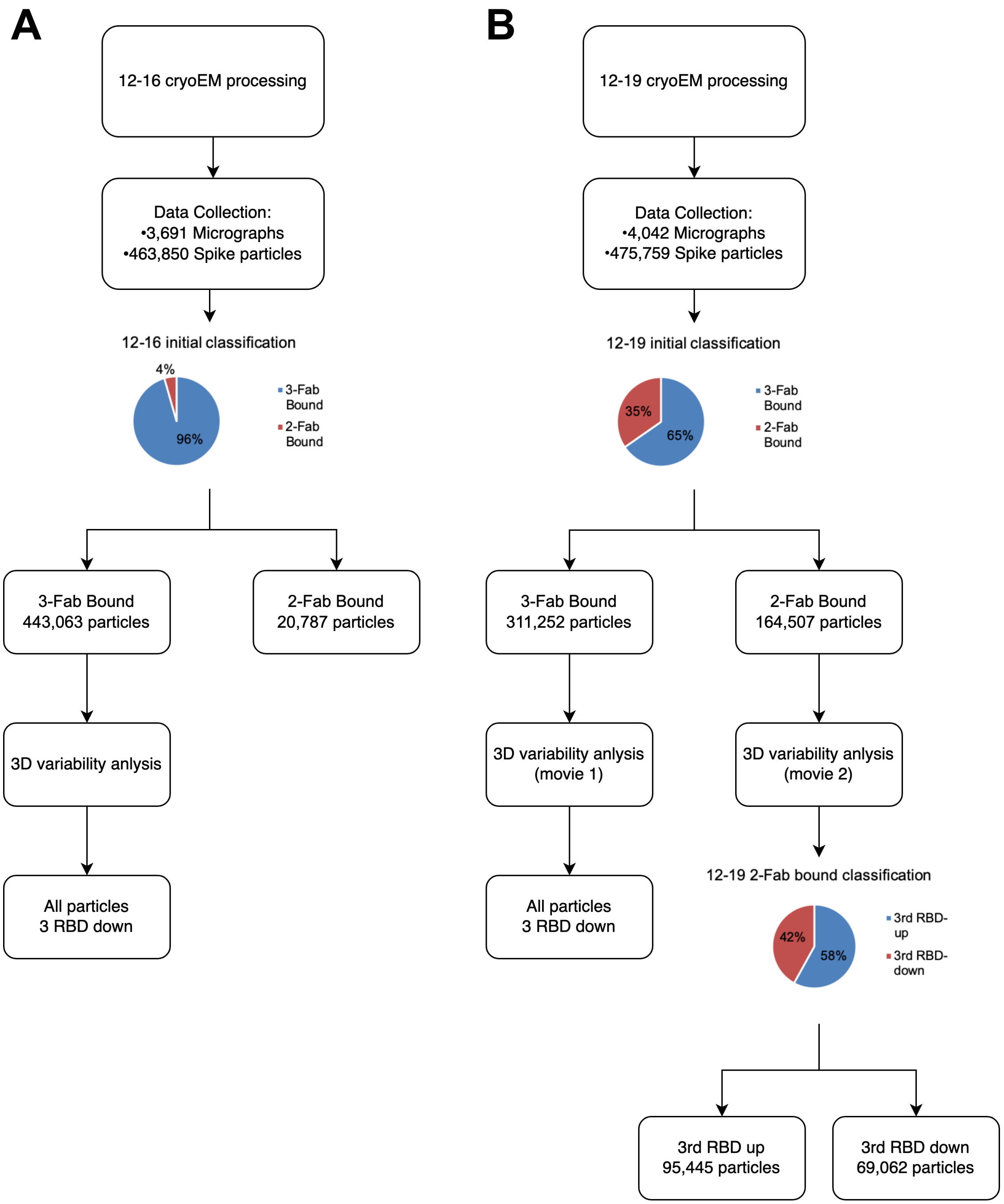

A
12-16 cryoEM processing
Data Collection:
•3,691 Micrographs
•463,850 Spike particles
12-16 initial classification
4%
96%
3-Fab Bound
2-Fab Bound
3-Fab Bound
443,063 particles
2-Fab Bound
20,787 particles
3D variability anlysis
All particles
3 RBD down
B
12-19 cryoEM processing
Data Collection:
•4,042 Micrographs
•475,759 Spike particles
12-19 initial classification
35%
65%
3-Fab Bound
2-Fab Bound
3-Fab Bound
311,252 particles
2-Fab Bound
164,507 particles
3D variability anlysis
(movie 1)
3D variability anlysis
(movie 2)
All particles
3 RBD down
12-19 2-Fab bound classification
42%
58%
3rd RBD-up
3rd RBD-down
3rd RBD up
95,445 particles
3rd RBD down
69,062 particles

### Figure S6

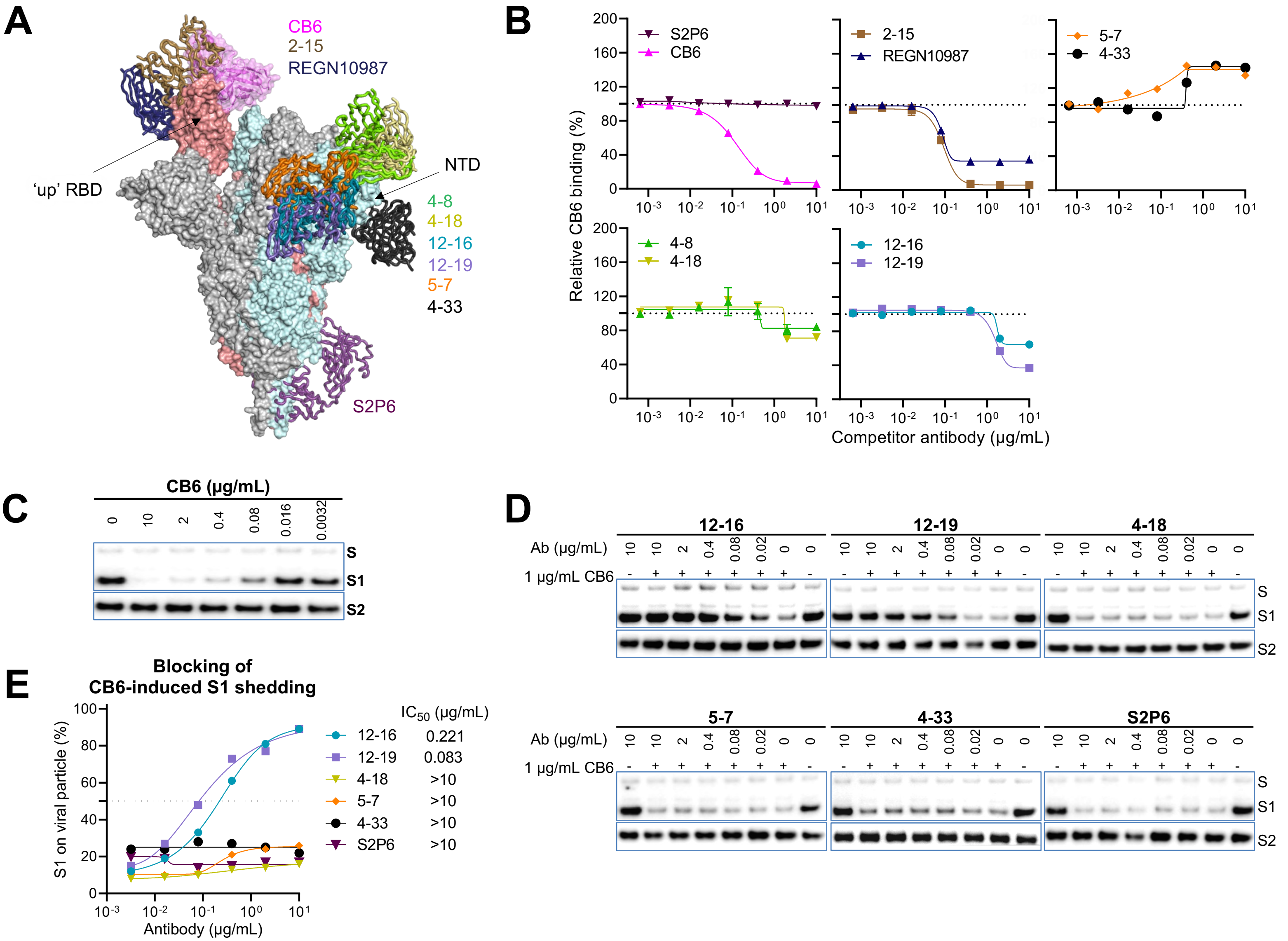
A
CB6
2-15
REGN10987
'up' RBD
NTD
4-8
4-18
12-16
12-19
5-7
4-33
S2P6
B
S2P6
CB6
2-15
REGN10987
5-7
4-33
4-8
4-18
12-16
12-19
Relative CB6 binding (%)
Competitor antibody (µg/mL)
C
CB6 (µg/mL)
0
10
2
0.4
0.08
0.016
0.0032
S
S1
S2
D
12-16
12-19
4-18
5-7
4-33
S2P6
Ab (µg/mL)
1 µg/mL CB6
S
S1
S2
E
Blocking of
CB6-induced S1 shedding
S1 on viral particle (%)
Antibody (µg/mL)
IC50 (µg/mL)
12-16 0.221
12-19 0.083
4-18 >10
5-7 >10
4-33 >10
S2P6 >10

### Figure S7

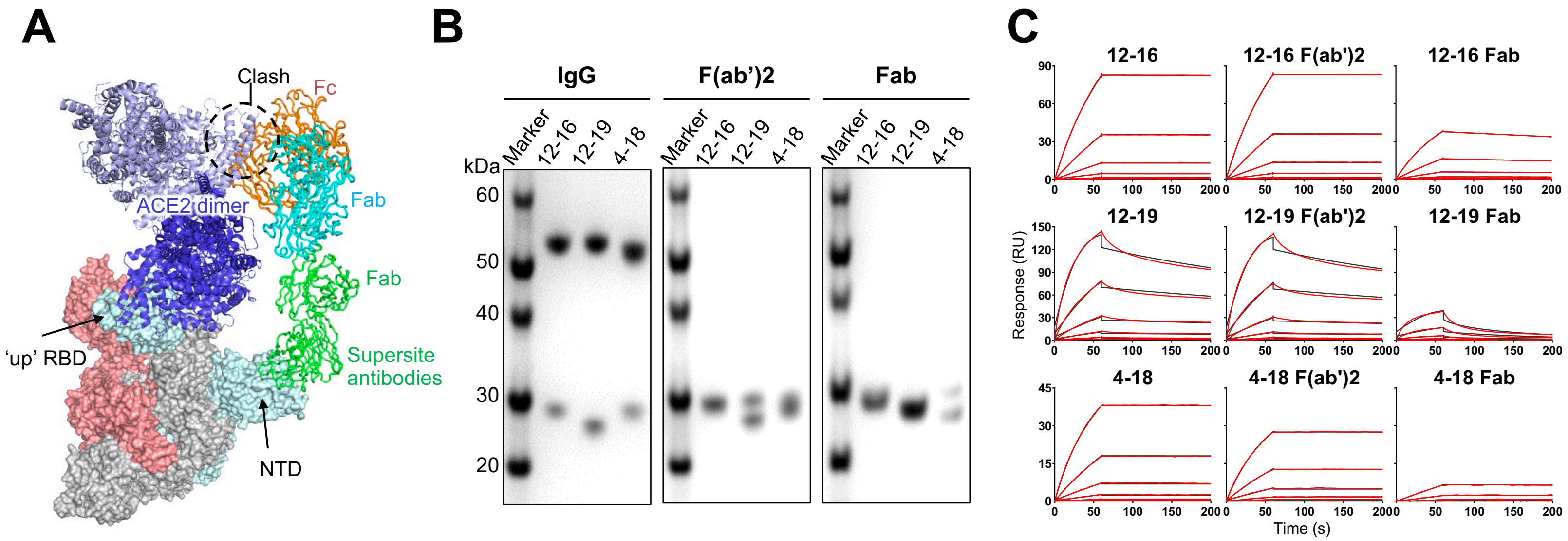

A
Clash
Fc
Fab
ACE2 dimer
Fab
'up' RBD
Supersite antibodies
NTD
B
IgG
F(ab')2
Fab
kDa
Marker
12-16
12-19
4-18
60
50
40
30
20
C
12-16
12-16 F(ab')2
12-16 Fab
12-19
12-19 F(ab')2
12-19 Fab
4-18
4-18 F(ab')2
4-18 Fab
Response (RU)
Time (s)

### Figure S8

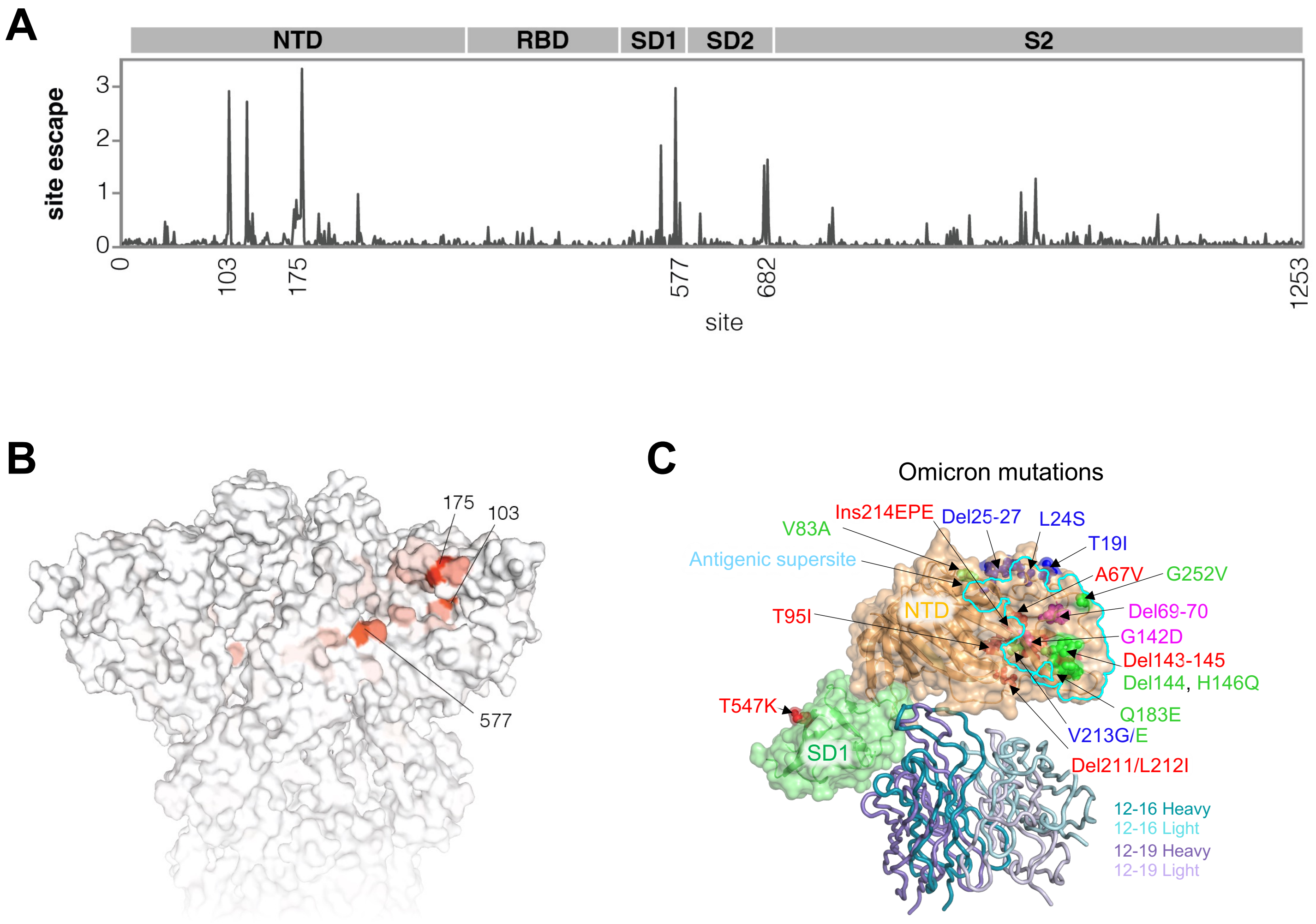
A
NTD
RBD
SD1
SD2
S2
site escape
0
1
2
3
0
103
175
577
682
1253
site
B
175
103
577
C
Omicron mutations
V83A
Ins214EPE
Del25-27
L24S
T19I
Antigenic supersite
A67V
G252V
NTD
Del69-70
T95I
G142D
Del143-145
Del144, H146Q
T547K
Q183E
V213G/E
SD1
Del211/L212I
12-16 Heavy
12-16 Light
12-19 Heavy
12-19 Light
